# Structural basis for PoxtA-mediated resistance to Phenicol and Oxazolidinone antibiotics

**DOI:** 10.1101/2021.06.18.448924

**Authors:** Caillan Crowe-McAuliffe, Victoriia Murina, Marje Kasari, Hiraku Takada, Kathryn Jane Turnbull, Yury S. Polikanov, Arnfinn Sundsfjord, Kristin Hegstad, Gemma C. Atkinson, Daniel N. Wilson, Vasili Hauryliuk

## Abstract

PoxtA and OptrA are ATP binding cassette (ABC) proteins of the F subtype (ABCF) that confer resistance to oxazolidinone, such as linezolid, and phenicol antibiotics, such as chloramphenicol. PoxtA/OptrA are often encoded on mobile genetic elements, facilitating their rapid spread amongst Gram-positive bacteria. These target protection proteins are thought to confer resistance by binding to the ribosome and dislodging the antibiotics from their binding sites. However, a structural basis for their mechanism of action has been lacking. Here we present cryo-electron microscopy structures of PoxtA in complex with the *Enterococcus faecalis* 70S ribosome at 2.9–3.1 Å, as well as the complete *E. faecalis* 70S ribosome at 2.2–2.5 Å. The structures reveal that PoxtA binds within the ribosomal E-site with its antibiotic resistance domain (ARD) extending towards the peptidyltransferase center (PTC) on the large ribosomal subunit. At its closest point, the ARD of PoxtA is still located >15 Å from the linezolid and chloramphenicol binding sites, suggesting that drug release is elicited indirectly. Instead, we observe that the ARD of PoxtA perturbs the CCA-end of the P-site tRNA causing it to shift by ∼4 Å out of the PTC, which correlates with a register shift of one amino acid for the attached nascent polypeptide chain. Given that linezolid and chloramphenicol are context-specific translation elongation inhibitors, we postulate that PoxtA/OptrA confer resistance to oxazolidinones and phenicols indirectly by perturbing the P-site tRNA and thereby altering the conformation of the attached nascent chain to disrupt the drug binding site.

## Introduction

Antibiotic resistance (ARE) is a growing threat to the efficacy of our current arsenal of clinically approved antimicrobial agents. The ATP-binding cassette (ABC) family of proteins are well-known for their role as multidrug resistance transporters, which use the energy of ATP hydrolysis to drive the efflux of antibiotics from the bacterial cytoplasm (Lubelski et al., 2007; Orelle et al., 2019). In recent years, it has become clear that a subfamily of ARE ABC proteins that belong to the subfamily F of the ABC family (ABCF) are not transporters – and thus do not confer resistance via efflux – but rather act via a direct target protection mechanism (Ero et al., 2021; Fostier et al., 2021; Murina et al., 2018; Murina et al., 2019; Ousalem et al., 2019; Sharkey et al., 2016; Sharkey and O’Neill, 2018; Wilson et al., 2020).

ARE-ABCF proteins confer resistance to a diverse range of antibiotics that inhibit protein synthesis by targeting the large subunit (LSU) of the ribosome. Based on the spectrum of antibiotic resistance that they confer ARE-ABCF proteins fall into three functional groups: (i) those that protect from pleuromutilins, lincosamides and streptogramins A (PLS_A_), (ii) those that protect from macrolides and streptogramin B (MS_B_), and, finally, (iii) those that protect from phenicols and oxazolidinones (PhO) (Ero et al., 2021; Fostier et al., 2021; Ousalem et al., 2019; Sharkey et al., 2016; Sharkey and O’Neill, 2018; Wilson et al., 2020). These functional groups do not map exactly to the phylogenetic tree of ARE-ABCFs, in which seven subclasses (ARE1-7) were originally distinguished, but are rather scattered amongst non-ARE ABCFs, implying that these resistance factors have arisen multiple times by convergent evolution (Murina et al., 2019). Despite the divergence in the spectrum of antibiotic resistance, the ARE-ABCFs share a common architecture and are comprised of two ABC nucleotide-binding domains (NBD1 and NBD2) that are separated by a helical linker, termed an ARD (antibiotic resistance domain), and, depending on the species, may have an additional “Arm” subdomain inserted within NBD1 as well as a C-terminal extension (CTE) (Murina et al., 2018). In fact, this architecture is similar to many non-ARE ABCFs, such as the *E. coli* housekeeping ABCF ATPase EttA, in which the ARD equivalent is shorter and referred to as a P-site tRNA-interaction motif (PtIM) (Boel et al., 2014; Chen et al., 2014), thus making it difficult to judge whether many ABCF proteins are actually resistance determinants or endogenous proteins of mostly unknown function (Murina et al., 2018).

Cryo-EM structures of ribosomes in complex with ARE-ABCFs that confer resistance to PLS_A_ (ARE1 VgaA_LC_ and VgaL, ARE2 VmlR, ARE3 LsaA) and MS_B_ (ARE1 MsrE) classes of antibiotics have revealed that these proteins bind within the E-site (Crowe-McAuliffe et al., 2018; Crowe-McAuliffe et al., 2021; Su et al., 2018), similar to that reported previously for the housekeeping non-ARE ABCF EttA (Chen et al., 2014). However, in contrast to EttA (Boel et al., 2014; Chen et al., 2014), the longer ARD of the ARE-ABCF proteins distorts the P-site tRNA, allowing the factor to access the peptidyl transferase center (PTC) on the LSU of the ribosome and dislodge the relevant antibiotics from their binding sites (Crowe-McAuliffe et al., 2018; Crowe-McAuliffe et al., 2021; Su et al., 2018). These structures revealed that there is often no steric overlap between the ARD of the ARE-ABCF and the drugs and even when there is a steric overlap, mutational analysis indicated that it is not strictly required for resistance (Crowe-McAuliffe et al., 2018; Crowe-McAuliffe et al., 2021; Su et al., 2018). Collectively, these results support a model where the ARE-ABCFs dislodge the drugs from the PTC by inducing a cascade of conformational changes within the 23S rRNA nucleotides that comprise the drug binding site (Crowe-McAuliffe et al., 2021). For MsrE, drug release was reported to occur in the presence of a non-hydrolysable ATP analog (Su et al., 2018), suggesting that ATP hydrolysis is not essential for drug release, but rather is needed for recycling of the factor from the ribosome.

In contrast to the relatively well-understood PLS_A_- and MS_B_-protecting ARE-ABCFs, mechanistic insights into how ARE7 OptrA and ARE8 PoxtA confer resistance to PhO antibiotics are lacking. The first ARE-ABCF from this group to be discovered is OptrA. This factor confers resistance to the phenicols, such as chloramphenicol and florfenicol, as well as the oxazolidinones linezolid and, to a lesser extent, tedizolid (Wang et al., 2015). OptrA was originally found on the conjugative plasmid pE349 from *Enterococcus faecalis* (Wang et al., 2015), but has subsequently been detected, both plasmid and chromosomal-encoded, across many Gram-positive enterococci, staphylococci and streptococci of human and animal origin (Cai et al., 2015; Fan et al., 2017; Lazaris et al., 2017; Li et al., 2016; Schwarz et al., 2021; Vorobieva et al., 2017; Wang et al., 2015). At least 69 variants of the *optrA* gene have been reported to date, differing by 1–20 aa substitutions, which corresponds to an amino acid identity of 97.1–99.8% compared to the first reported OptrA sequence (Schwarz et al., 2021). Moreover, evidence for horizontal transfer of OptrA to Gram-negative bacteria, such as *Campylobacter coli*, has also been recently described (Liu et al., 2020; Tang et al., 2020). While oxazolidinones are not clinically efficient against Gram-negative pathogens, this raises the concern of possible co-selection of antibiotic co-resistance, i.e. selection for simultaneous transfer and spread of several antibiotic resistance genes encoded by one mobile genetic element. In addition to OptrA, a second ARE-ABCF from this group was detected by bioinformatic analysis of the genome of a methicillin-resistant *Staphylococcus aureus* (MRSA) strain AOUC-0915 isolated from a cystic fibrosis patient at the Florence Careggi University Hospital in Florence, Italy (Antonelli et al., 2018). Expression of the resistance determinant in *S. aureus, E. faecalis* and *E. coli* was reported to confer resistance to <u>p</u>henicol-<u>ox</u>azolidinone-tetracycline antibiotics, and was therefore termed PoxtA (Antonelli et al., 2018). To date, PoxtA has been found exclusively in *Enterococcus* and *Staphylococcus* species, most frequently in *E. faecium* isolates of both human and animal origin (Schwarz et al., 2021).

In the absence of structures of OptrA and PoxtA on the ribosome, it has remained unclear how these ARE-ABCFs confer antibiotic resistance. Both chloramphenicol and linezolid bind at the PTC and inhibit the elongation phase of protein synthesis (Wilson, 2014). However, their activity is context-specific such that translation arrest is most efficient when the nascent polypeptide chain on the ribosome carries an alanine residue and, to a lesser extent, serine or threonine in its penultimate position (Choi et al., 2020; Marks et al., 2016; Vazquez-Laslop and Mankin, 2018). Although the ARDs of OptrA and PoxtA are slightly longer (4–5 aa) than the PtIM of non-ARE ABCFs such as EttA, they are considerably shorter than the ARDs of ARE-ABCFs from other groups, and at least 20 amino acids shorter than other ARE-ABCFs for which structures have been reported (**Fig. 1**). Thus, assuming OptrA and PoxtA bind similarly to the ribosome as other ARE-ABCFs, the ARDs are unlikely to be able to reach into the PTC to dislodge the drugs from their binding site (Wilson et al., 2020). Moreover, it is hard to rationalize how PoxtA also confers resistance to tetracycline antibiotics, which bind near the decoding site on the small subunit (SSU), which is located far from the PTC on the LSU (Antonelli et al., 2018).

**Fig. 1.**
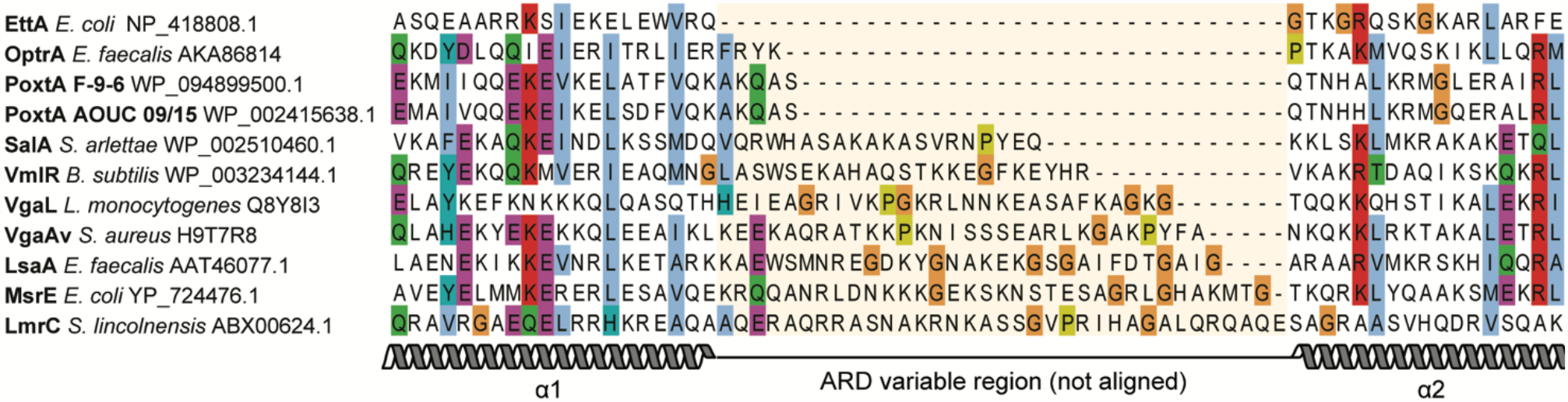
Alignment of the ARDs from diverse bacterial ABCFs. The PoxtA and OptrA ARDs are slightly longer (4–5 amino acids) than the equivalent region in EttA, but significantly shorter than for other ARE-ABCFs. The central region with orange highlight, which includes the ARD loop and some of the adjacent helices in some proteins, was not aligned but simply ordered by length. Sequences were aligned with MAFFT (Katoh and Standley, 2013) and edited by hand to reduce gap placement.

Here we have systematically characterized the PoxtA and OptrA resistance determinants, revealing that both increase the minimum inhibitory concentrations (MICs) to phenicols, such as chloramphenicol, thiamphenicol and florfenicol, as well as to the oxazolidinone linezolid, but not to macrolides, pleuromutilins, lincomycins, streptogramins that also bind at the PTC. Morever, we find no evidence for either PoxtA or OptrA confering resistance to non-PTC binding antibiotics, such as tetracycline. Cryo-EM structures of PoxtA on the ribosome reveal that it binds in the E site and, despite the short ARD, still induces a distortion of the P-site tRNA, leading to retraction of its CCA-end from the PTC. Unlike for other ARE-ABCFs, we observe no conformational rearrangements within the 23S rRNA around the drug binding sites within the A site of the PTC upon binding of PoxtA to the ribosome. This leads us to propose a model whereby the distortion of the P-site tRNA by PoxtA (and OptrA) reduces the affinity of the drugs for their binding site by altering the context and therefore interaction of the nascent polypeptide chain with respect to the drugs.

## Results

### PoxtA and OptrA confer resistance to phenicols and oxazolidinones, but not tetracyclines

To systematically characterise the antibiotic resistance profiles of PoxtA and OptrA, representatives of these ARE-ABCF groups were expressed in an *E. faecalis* TX5332 strain where the *lsaA* gene had been disrupted (Δ*lsaA*), and MICs were determined for phenicol (chloramphenicol, thiamphenicol and florfenicol), oxazolidinone (linezolid), macrolide (erythromycin, azithromycin and leucomycin), lincosamide (lincomycin and clindamycin), pleuromutilin (tiamulin, retapamulin), streptogramin A and B (Virginiamycin M1 and S1, respectively) and tetracycline antibiotics (**Table 1**). We have characterised OptrA E35048 from *E. faecium* (Morroni et al., 2018), OptrA ST16 from the clinical *E. faecalis* ST16 isolate (Vorobieva et al., 2017), PoxtA AOUC-0915 from MRSA (Antonelli et al., 2018) and, finally, PoxtA E9F6 from a multidrug-resistant ST872 *E. faecium* clinical isolate 9-F-6 (Sivertsen et al., 2018). As controls, we determined MICs for the *E. faecalis* Δ*lsaA* strain transformed with the empty vector plasmid pCIE_spec_, as well as expressing LsaA, the native genome-encoded ARE-ABCF from *E. faecalis*.

**Table 1.** Antibiotic resistance spectra LsaA, OptrA and PoxtA ARE-ABCFs.

|  | Minimum Inhibitory Concentration (MIC, µg/mL) |  |  |  |  |  |
| --- | --- | --- | --- | --- | --- | --- |
|  | pCIE<sub>spec</sub><br>vector | LsaA | OptrA<br>ST16 | OptrA<br>35048 | PoxA<br>EF9F6 | PoxA<br>AOUC 0915 |
| Chloramphenicol | 2-4 | 2-4 | **8-16** | **4-8** | **4-8** | **4-8** |
| Thiamphenicol | 4 | 4 | **32-64** | **16-32** | **8-16** | **32-64** |
| Florfenicol | 1-2 | 1-2 | **16-32** | **8** | **2-4** | **16-32** |
| Linezolid | 1 | 1 | **8** | **2-4** | **2-4** | **4-8** |
| Erythromycin | 0.5-1 | 0.5 | 0.5 | 0.5 | 0.5 | 0.5 |
| Azithromycin | 0.5 | 0.5 | 0.5 | 0.5 | 0.5 | 0.5 |
| Leucomycin | 0.5 | 0.5 | 0.5 | 0.5 | 0.5 | 0.5 |
| Lincomycin | 0.125 | **16-32** | 0.125 | 0.125 | 0.125 | 0.125 |
| Clindamycin | 0.0156 | **16** | 0.0156 | 0.0156 | 0.0156 | 0.0156 |
| Tiamulin | 0.0625 | **128** | 0.0156 | 0.0156 | 0.031 | 0.031 |
| Retapamulin | 0.0156 | **>64** | 0.0156 | 0.0156 | 0.0156 | 0.0156 |
| Virginiamycin M1 | 4 | **>128** | 4 | 4 | 4 | 4 |
| Virginiamycin S1 | 8 | 8 | 8 | 8 | 8 | 8 |
| Tetracycline | 0.25 | 0.25 | 0.25 | 0.25 | 0.25 | 0.25 |

BHI media supplemented with 2 mg/mL kanamycin (to prevent *Isa* revertants), 0.1 mg/mL spectinomycin (to maintain the pCIE<sub>spec</sub> plasmid), 100 ng/mL of cCF10 peptide (to induce expression of ABCF proteins) as well as increasing concentrations of antibiotics was inoculated with 5 x 10<sup>5</sup> CFU/mL (≈OD<sub>600</sub> 0.0005) of *E. faecalis* Δ*IsaA* (*Isa*::Kan) strain TX5332 transformed either with empty pCIE<sub>spec</sub> plasmid, or with pCIE<sub>spec</sub> derivatives for expression of ARE-ABCF proteins. After 16-20 hours at 37 °C without shaking, the presence or absence of bacterial growth was scored by eye. The MIC values that are higher than the empty vector control are shown in bold. The experiments were performed in triplicates.

In agreement with earlier reports, expression of LsaA confers resistance to PLS_A_ antibiotics, but not PhO, MS_B_ or tetracycline (Crowe-McAuliffe et al., 2021; Singh et al., 2002). By contrast, cCF10-inducible expression of either OptrA or PoxtA results in from 2- to 16-fold MIC increase for PhO antibiotics, and does not, as expected, result in resistance against either PLS_A_ or MS_B_, as observed in earlier reports (Antonelli et al., 2018; Wang et al., 2015). While the earlier study (Antonelli et al., 2018) reported a minor protective effect of PoxtA against tetracycline (a two-fold increase in MIC for *E. faecalis* and *S. aureus*), in our hands expression of neither of the PoxtA variants resulted in any increase in MIC for this antibiotic. Notably, the lack of effect of PoxtA AOUC-0915 expression on the MIC for the tetracycline antibiotic tigecycline in either *E. faecalis* or *S. aureus* (Antonelli et al., 2018) is consistent with the original *E. faecium* 9-F-6 strain from which we have isolated PoxtA E9F6 also being susceptible to tigecycline (**Table S1**). Therefore, we concluded that the antibiotic resistance spectrum of PoxtA is similar, if not identical, to that of OptrA. This similarity appears to be a case of convergent evolution, as there is no phylogenetic support for PoxtA and OptrA being more closely related to each other than to any other ABCF subfamily (**Fig. S1**). Indeed, OptrA is more closely related to the vertically inherited and probable housekeeping ABCF YdiF of Firmicutes (84% bootstrap support, **Fig. S1**), while the relationship of PoxtA to other subfamilies is unresolved (bootstrap support below 50%). On these grounds we conclude that PoxtA-like proteins constitute a separate ARE subfamily which we call ARE8.

### Cryo-EM structures of PoxtA-70S complexes

Our attempts to reconstitute OptrA- and PoxtA-70S ribosome complexes *in vitro* were unsuccessful due to problems obtaining soluble homogenous OptrA and PoxtA proteins. Therefore, we employed an *in vivo* pull-out approach with strains overexpressing C-terminally FLAG_3_-tagged OptrA and PoxtA proteins, as used recently to generate other ARE-ABCF–ribosome complexes (Crowe-McAuliffe et al., 2021). We note that the inclusion of chloramphenicol or linezolid blocks OptrA/PoxtA overexpression and therefore complex formation and purification was performed in the absence of the antibiotic. We expressed both the wild-type ATPase-competent OptrA- and PoxtA and the ATPase-deficient variants bearing Glu-to-Gln substitutions in both NBD cassettes (EQ_2_). Such EQ_2_ variants have been successfully employed to trap other ABCF proteins on the ribosome because they allow binding but prevent hydrolysis of ATP (Chen et al., 2014; Crowe-McAuliffe et al., 2018; Crowe-McAuliffe et al., 2021; Kasari et al., 2019). The OptrA-EQ_2_ variant was earlier shown to have a compromised *in vitro* ATPase activity and be incompetent in promoting antibotic resistance *in vivo* (Zhong et al., 2018). Affinity purification via the FLAG_3_ tag was performed in the presence of 0.5 mM ATP from clarified lysates prepared from the *E. faecalis* Δ*lsaA* strain expressing the FLAG_3_-tagged ARE-ABCF, either wildtype or EQ_2_-variant. Analysis of the elution fractions from the purifications indicated that only the *E. faecium* PoxtA(E9F6)-EQ_2_ factor was bound stably to the ribosome (**Fig. S2**). Furthermore, our attempts with OptrA carrying a single individual EQ substitution (E470Q) were equally unsuccessful. Therefore, we focused on the PotxA(E9F6)-EQ_2_–70S sample and subjected it to structural analysis using single-particle cryo-EM.

Using a Titan Krios transmission electron microscope with a K2 direct electron detector, we collected 3,640 micrographs which, after 2D classification, yielded 140,310 ribosomal particles (**Fig. S3**). *In silico* sorting revealed that 80% of these particles contained an additional density for PoxtA and/or tRNAs, which after 3D refinement resulted in a cryo-EM map of *E. faecalis* 70S ribosome with an average resolution of 2.4 Å (**Figure S4A**). Subsequent mutibody refinement yielded average resolutions of 2.2 Å and 2.5 Å for the LSU and SSU, respectively (**Fig. S4B–E**). The increase in resolution compared to the previous *E. faecalis* 70S ribosome models at 2.8–2.9 Å (Crowe-McAuliffe et al., 2021; Murphy et al., 2020) is evident from improved quality and features of the cryo-EM density, including visualization of some rRNA modifications (e.g. N2-methylguanosine), water molecules and hydrated magnesium ions (**Fig. S5A–F, Table S2**).

Further subsorting of ribosomal particles using a mask focused on the intersubunit space yielded four defined classes, which we refer to as states I–IV (**Fig. S3**). States I and II contained density for PoxtA bound in the E site and tRNA in the P site, and had average resolutions of 2.9 Å and 3.0 Å, respectively (**Fig. 2A** and **Fig. S6A–B, Table S2**). State II differed from state I by only a slight rotation of the SSU relative to the LSU. State III was similar to state II, but additionally contained an A-site tRNA (**Fig. 2B**), whereas state IV contained P-site tRNA only (**Fig. 2C**), presumably because PoxtA dissociated during sample preparation. States III and IV were also refined, resulting in final reconstructions with average resolutions of 2.9 Å and 3.1 Å, respectively (**Fig. S6A–B, Table S2**). The cryo-EM density for PoxtA in states I–III was generally well-resolved (**Fig. S6C**), enabling a reliable model to be built for NBD1, NBD2 and the ARD (**Fig. 2D**). By contrast, the Arm domain, which interacts with uL1, appeared flexible (**Fig. S6C**) and could only be modeled as a rigid body fit of the two α-helices (**Fig. 2D**). The best-resolved region of PoxtA was the ARD, consisting of two α-helices (α1 and α2) and the ARD loop (**Fig. 2D** and **Fig. S6C**), where the majority of sidechains could be modeled unambiguously (**Fig. 2E**). Additional density located between NBD1 and NBD2 of PoxtA was attributed to two ATP molecules (ATP-1 and ATP-2) and a magnesium ion (**Fig. 2F**), as expected from the use of the ATPase-deficient PoxtA-EQ_2_ variant. As observed for other ribosome-bound ARE-ABCF structures (Crowe-McAuliffe et al., 2018; Crowe-McAuliffe et al., 2021; Su et al., 2018), the NBDs of PoxtA adopt a closed conformation, which is also consistent with the inability to hydrolyze ATP. In all states, the anticodon stem loop (ASL) and acceptor arm, including the CCA-end, are well-resolved, whereas the elbow region of the tRNAs exhibit some flexibility (**Fig. S6D**). The density for the P-site tRNA is consistent with initiator tRNA^fMet^, indicating that in the absence of chloramphenicol- or linezolid-stalled ribosomes, the PoxtA-EQ_2_ variants bind to the vacant E site of initiation complexes, as observed previously for other ARE-ABCF-ribosome complexes (Crowe-McAuliffe et al., 2021). In state IV, clear density for the fMet moiety attached to the P-site tRNA is evident, whereas state III appears to be a post-peptide bond formation state with a deacylated P-site tRNA and A-site tRNA bearing a dipeptide. In states I and II, which contain PoxtA but lack A-site tRNA, some density for the fMet moiety on the distorted P-site tRNA is evident but is poorly resolved.

**Fig. 2.**
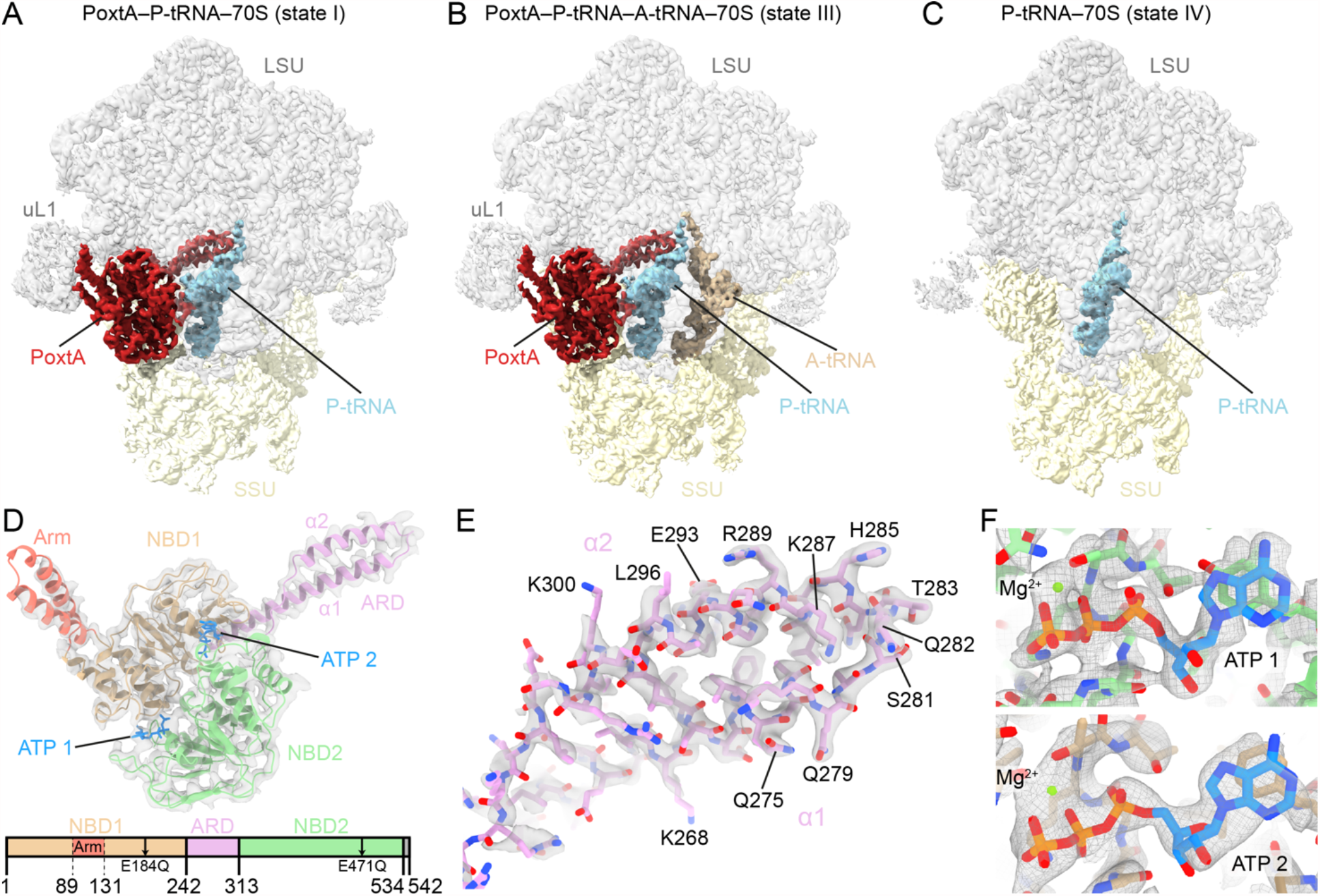
Cryo-EM structures of PoxtA–70S complexes. (*A*–*C*) Cryo-EM maps with isolated densities for (*A, B*) *E. faecium* PoxtA (red) in complex with the *E. faecalis* 70S ribosome and (*A*) P-tRNA (cyan) or (*B*) P-tRNA (cyan) and A-tRNA (tan), (*C*) P-tRNA (cyan) only, with small subunit (SSU, yellow) and large subunit (LSU, grey). (*D*) Density (grey mesh) with molecular model of PoxtA from (*A*) coloured according to domain as represented in the associated schematics: nucleotide binding domain 1 (NBD1, tan), antibiotic-resistance domain (ARD, pink), nucleotide binding domain 2 (NBD2, green) and C-terminal extension (CTE, grey, not modelled). α1 and α2 indicate the two α-helices of the ARD interdomain linker. In (*D*), the ATP nucleotides are coloured blue. (*E*) Close view of the ARD tip from state I with sharpened map. (*F*) Close view of ATPs bound by PoxtA (state I).

### Interaction of PoxtA with the ribosome and P-site tRNA

In states I–III, PoxtA is located within the ribosomal E site (**Fig. 3A**) and generally binds similarly to that observed for other ARE-ABCF proteins, such as VmlR, MsrE, LsaA, VgaL and VgaA_LC_ (Crowe-McAuliffe et al., 2018; Crowe-McAuliffe et al., 2021; Su et al., 2018), as well as the non-ARE-ABCF protein EttA (Chen et al., 2014). Unlike these ARE-ABCFs that lack an Arm subdomain (or have a short Arm in the case of LsaA) (Murina et al., 2018), the Arm subdomain of PoxtA (and OptrA (Murina et al., 2018)) is prominent, like that of EttA (Boel et al., 2014; Chen et al., 2014), and stabilises an open conformation of the L1 stalk via direct interaction with domain II of uL1 (**Fig. 3A**). Additional contacts to the 23S rRNA helices H77/H78 of the L1 stalk are evident from the NBD1 of PoxtA, as are interactions for NBD1 with H68 and bL33 on the LSU (**Fig. 3A**). By contrast, NBD2 of PoxtA spans across the intersubunit interface, establishing interactions with uL5 on the LSU as well as uS7 and h41 on the SSU (**Fig. 3A**). NBD2 also interacts directly with the elbow region of the P-site tRNA, namely, with the G19-C56 basepair that links the D- and T-loops (**Fig. 3A**,**B**). Here, Ser430 of PoxtA is within hydrogen bonding distance of the N7 of G19 and the sidechain of Arg426 of PoxtA stacks upon the nucleobase of C56 of the T-loop (**Fig. 3B**). However, it is the ARD that makes the most extensive interactions with the P-site tRNA, establishing a complex network of hydrogen bonding interactions with the acceptor arm and CCA-end (**Fig. 3A**). In particular, two glutamine residues, Gln275 and Gln279, located at the distal end of the α-helix α1 of the ARD insert into the minor groove of the acceptor arm where hydrogen bond interactions can form with the C3-G70 base-pair (**Fig. 3C**). Hydrogen bonding is also possible from the ε-amino group of Lys278 and the backbone carbonyl of Gln275 of PoxtA with the phosphate-oxygen of G4 and the ribose-oxygen of C71 of the P-tRNA, respectively. The loop region of the ARD of PoxtA interacts predominantly with the single-stranded CCA-3’ end of the P-site tRNA (**Fig. 3D**). Ser281 is within H-bonding distance to the phosphate-oxygen of A73, whereas the sidechain of Gln282 stacks upon the base of C74 and can interact with the ribose-hydroxyl of A72 (**Fig. 3D**). C75 of the P-site tRNA is also stabilized by indirect contacts with the backbone carbonyl of Thr283 via a water molecule, as well as a direct H-bond with the sidechain of His285, the first residue of α-helix α2 of the ARD of PoxtA (**Fig. 3E**). The ARD is stabilized by multiple contacts between residues within α-helix α2 and rRNA nucleotides located in H74 and H93. For example, the sidechains of Arg294 and Arg297 contact nucleotides A2595–G2598 (*E. coli* numbering used throughout) located within the loop of H93 of the 23S rRNA, and Glu293 interacts with G2597 directly as well as G2598 via a putative water molecule (**Fig. 3F**).

**Fig. 3.**
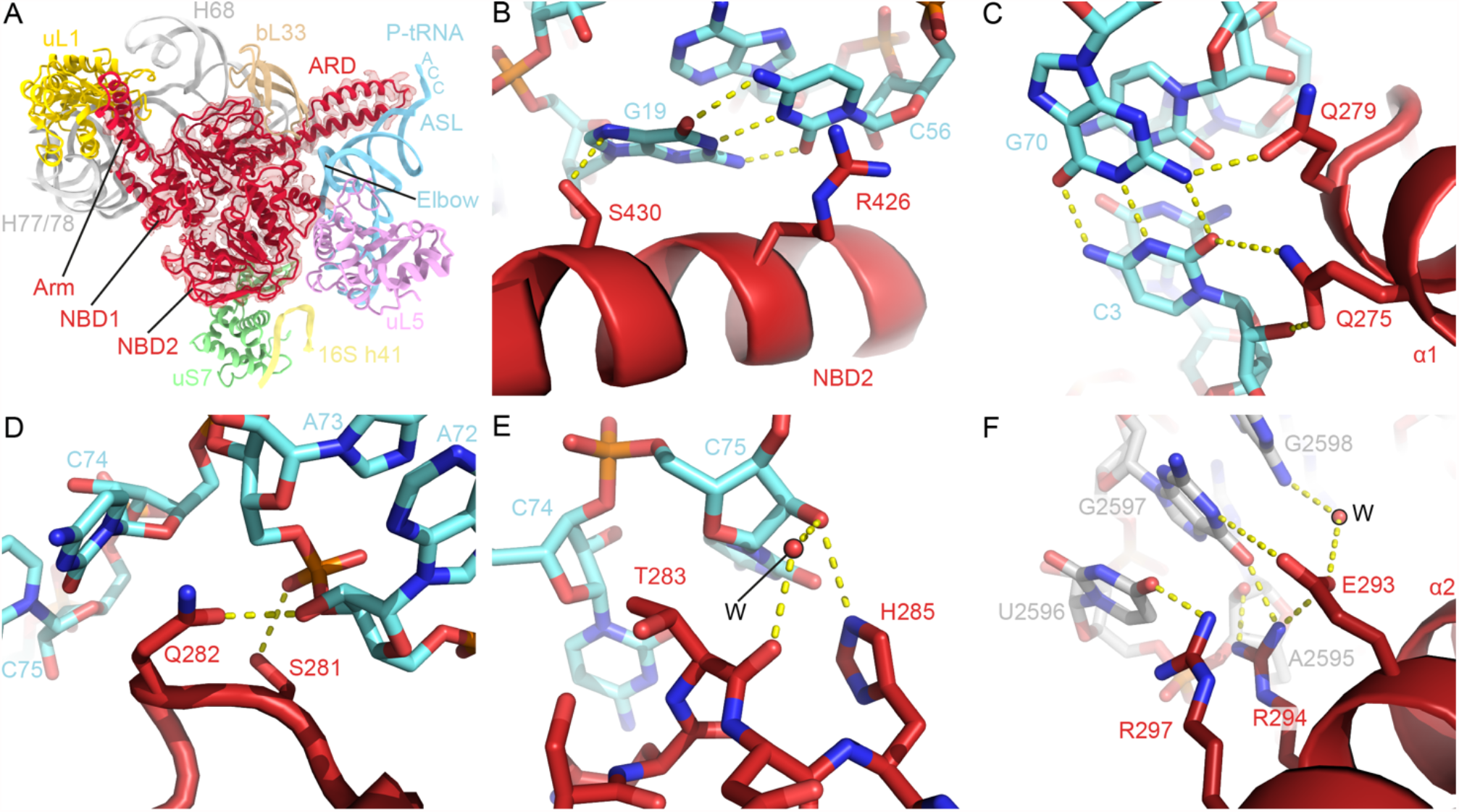
Interactions between PoxtA and the ribosome-P-tRNA complex. (*A*) Overview of PoxtA interactions with the 23S rRNA (grey), 16S rRNA h41 (yellow), uL1 (gold), uS7 (green), uL5 (pink), bL33 (tan) and the P-tRNA (light blue). (*B*–*E*) Interactions between the P-tRNA elbow (light blue) and the PoxtA NBD2 (*B*), the P-tRNA acceptor stem (light blue) and the PoxtA ARD (*C*), the ARD and the CCA end (*D, E*). (*F*) An interaction between the PoxtA ARD α2 and the 23S rRNA. The high-resolution model from the combined 70S volume was used.

### PoxtA perturbs the position of the CCA-end of the P-site tRNA at the PTC

Despite the short ARD, binding of PoxtA to the E site nevertheless causes a distortion of the P-site tRNA when compared to the canonical P-site tRNA binding position such as that observed in state IV (**Fig. 4A, B**). While the ASL remains fixed in position on the SSU where it decodes the P-site codon of the mRNA, the elbow region shifts towards the E site by ∼6-7 Å, thus bringing it into contact with NBD2 of PoxtA (**Fig. 4B**). The shift of the elbow region is very similar to that observed for the distorted P-site tRNAs observed on the ribosome in the presence of the other ARE-ABCFs (Crowe-McAuliffe et al., 2018; Crowe-McAuliffe et al., 2021; Cue et al., 2009; Su et al., 2018), such as LsaA (**Fig. 4C**). However, in the PoxtA-70S complex, the distortion is accompanied by a smaller ∼4 Å shift of the acceptor arm of the P-site tRNA away from the PTC (**Fig. 4D**), whereas for other ARE-ABCF complexes, the CCA-end of the P-site tRNA completely vacates the PTC due to the presence of the longer ARD (as illustrated here for LsaA in **Fig. 4C**). Although PoxtA contains a shorter ARD than other ARE-ABCFs, the loop of the ARD still contacts the acceptor stem of the P-tRNA, precluding canonical interactions between the tRNA and the ribosome (**Fig. 4D**). As a consequence, the single-stranded CCA-3’ end of the P-site tRNA becomes contorted in the presence of PoxtA and the canonical interactions of the C75 and C74 of a P-site tRNA with 23S nucleotides G2251 and G2252, respectively, of the P-loop (H80) are disrupted (**Fig. 4E, F**). This results in a shift in register such that C75 basepairs with G2252 and C74 stacks upon Gln282 of PoxtA and interacts with G2253 and C2254 (**Fig. 4F**). The register shift is reminiscent of, but distinct to, that observed for the P_int_-tRNA in *E. coli* 70S ribosome complexes formed in the presence of the antimicrobial peptide apidaecin and the termination release factor RF3 (Graf et al., 2018).

**Fig. 4.**
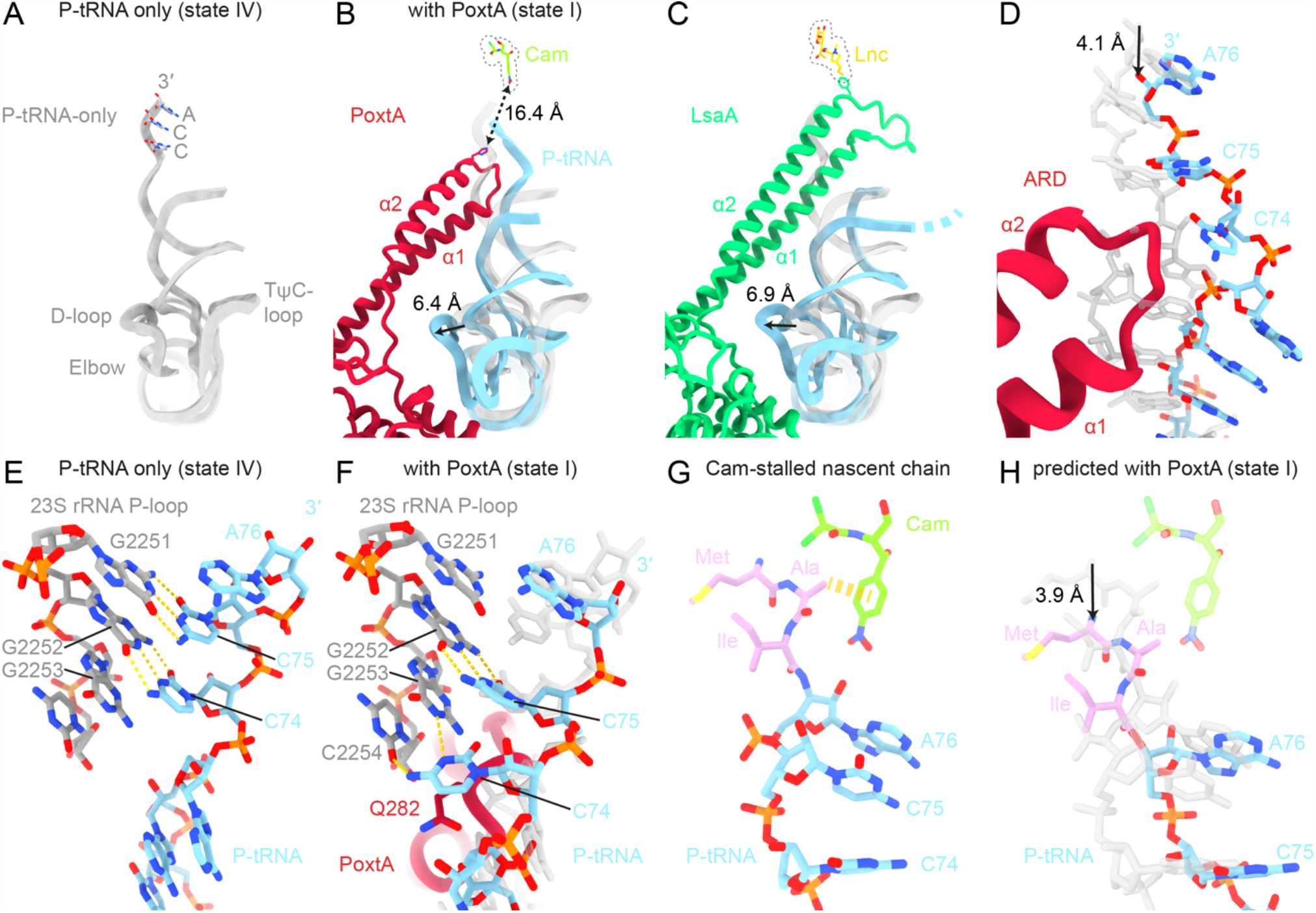
Conformational changes at the peptidyltransferase centre. (*A, B*) Comparison of position of CCA-end of the P-site tRNA at PTC from the (*A*) *E. faecalis* 70S-P-tRNA only complex (grey), (*B*) PoxtA-70S complex (state I, PoxtA in red and P-tRNA in light blue), and (*C*) LsaA-70S complex (PDB ID 7NHK, (Crowe-McAuliffe et al., 2021), LsaA in green and P-tRNA in light blue). Models of chloramphenicol for PoxtA (PDB ID 6ND5, (Svetlov et al., 2019)) and lincomycin for LsaA (PDB ID 5HKV) (Matzov et al., 2017) are superimposed for referece (*D*) Close-up of (*B*) showing P-tRNA acceptor stem distortion induced by PoxtA binding. (*E, F*) Interaction between the 23S rRNA P-loop (grey) and P-tRNA (light blue) for (*E*) the P-tRNA-only complex (state IV) and (*F*) with bound PoxtA (state IV P-tRNA is overlayed in transparent grey for comparison). (*G, H*) Indirect modulation of the chloramphenicol binding site by PoxtA. (*G*) The structure of chloramphenicol (cam) stabilised by a nascent peptide chain with alanine in the −1 position (Syroegin et al., 2021). (*H*) modelled shift of the nascent chain shown in (*G*) induced by PoxtA displacement of the P-tRNA CCA end. The P-tRNA from (*G*) is overlayed in transparent grey for comparison.

A comparison of the binding position of the ARD of PoxtA with that of the phenicol (chloramphenicol) and oxazolidinone (linezolid) antibiotics reveals that there is no steric overlap between PoxtA and either drug (**Fig. 4B**). Indeed, His285 of PoxtA, which comes closest to the drugs, is still located >16 Å away, and furthermore the ARD of PoxtA is partitioned from the drugs by the CCA-3’ end of the P-site tRNA (**Fig. 4B**). This raises the question as to whether PoxtA dislodges these drugs from the ribosome using an indirect mechanism, such as by inducing conformational changes within the 23S rRNA nucleotides comprising the drug binding site(s), as proposed for other ARE-ABCFs (Crowe-McAuliffe et al., 2018; Crowe-McAuliffe et al., 2021; Su et al., 2018). To examine this, we compared the conformation of the PTC nucleotides comprising the drug binding sites in our structures with those in the presence of chloramphenicol and linezolid. We could find no evidence that the presence of PoxtA induces any conformational changes within the drug binding sites that would lead to dissociation of the drugs (**Fig. S7–S8**). In fact, we note that the conformation of the A-site pocket where the drugs bind is very similar, if not identical, for states I–IV, regardless of whether PoxtA is present (state I–III) or absent (state IV), or whether the A-site is vacant (state I–II, IV) or occupied (state III) (**Fig. S7–S8**). Although this suggests that PoxtA does not dislodge the drugs from the ribosome by altering the 23S rRNA in the binding site, we cannot exclude that such conformational changes occur upon ATP hydrolysis or upon dissociation of PoxtA from the ribosome.

An alternative scenario is that the distortion of the P-site tRNA induced by PoxtA binding indirectly reduces the affinity of the drugs for their A-site binding pocket. Both chloramphenicol and linezolid are context-specific inhibitors of translation elongation, such that the highest translation arrest activity occurs when the penultimate amino acid (−1 position) attached to the CCA-end of the P-site tRNA is alanine, and to a lesser extent, serine or threonine (Choi et al., 2020; Marks et al., 2016; Vazquez-Laslop and Mankin, 2018). Structures of chloramphenicol with peptidyl-tRNA mimics reveal an intimate interaction between the drug and the nascent polypeptide chain, illustrating how alanine in the −1 position stabilizes drug binding via a CH-π interaction (**Fig. 4G**) (Syroegin et al., 2021). Since the distortion of the P-site tRNA by PoxtA involves a shift out of the PTC by 4 Å, effectively altering the nascent chain register by one amino acid, this would also result in a shift of the alanine away from chloramphenicol (**Fig. 4H**) and thereby perturb the interactions and reduce the affinity of the drug for the ribosome.

## Discussion

### Model of antibiotic resistance mediated by PoxtA and OptrA

Based on the structures of the PoxtA-ribosome complexes determined here, as well as the available literature on the mechanism of action of oxazolidinones and phenicols, we propose a model for how PoxtA can confer antibiotic resistance to these antibiotic classes (**Fig. 5**). As mentioned, linezolid and chloramphenicol are context-specific inhibitors that stall the ribosome during translation elongation with the peptidyl-tRNA in the P site and the drug bound within the A-site pocket of the PTC (**Fig. 5A**) (Choi et al., 2020; Marks et al., 2016; Vazquez-Laslop and Mankin, 2018). The amino acid in the −1 position of the nascent polypeptide chain influences the strength of the arrest. Specifically, alanine – and, to a lesser extent, serine and threonine – elicit the strongest arrest (Choi et al., 2020; Marks et al., 2016; Vazquez-Laslop and Mankin, 2018), apparently due to direct interaction with the ribosome-bound drug (**Fig. 5A**) (Syroegin et al., 2021).

**Fig. 5.**
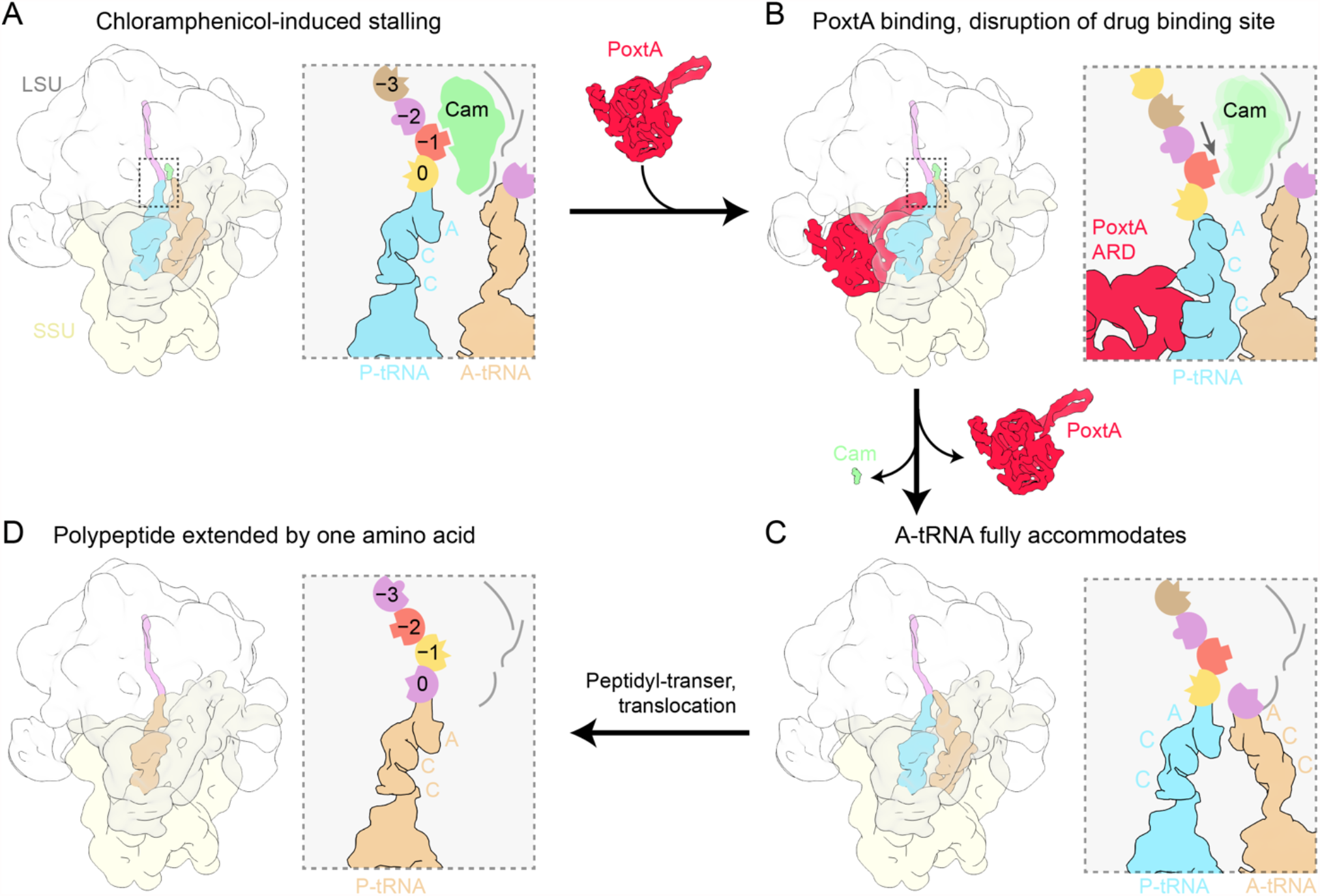
Model for ribosome protection by PoxtA. (*A*) Elongating ribosomes are stalled by PhO antibiotics with a peptidyl-tRNA in the P-site. The side chain of the amino acid at position −1 contributes to the PhO binding site. (*B*) Stalled ribosomes are recognized by PoxtA, which induces a shifted P-site tRNA and nascent chain, thereby disrupting the PhO binding site. (*C*) After dissociation of the PhO drug and PoxtA, the P-tRNA and nascent chain return to the regular conformation. (*D*) Accommodation of an A-tRNA occludes the drug binding site. After peptidyl transfer and translocation, amino acid at position −1 would change, thereby resetting the PhO binding site.

We propose that PoxtA recognizes these drug-arrested ribosomes and binds to the vacant E site (**Fig. 5B**). In contrast to other ARE-ABCFs where the CCA end is completely removed from the PTC (Crowe-McAuliffe et al., 2018; Crowe-McAuliffe et al., 2021; Su et al., 2018), the binding of PoxtA causes the CCA-end to shift by only ∼4 Å. Although modest, such a shift would be sufficient to change the register of the nascent polypeptide chain with respect to the drug, such that the drug can no longer interact with the amino acid in the −1 position, thereby decreasing its affinity for the ribosome and leading to its dissociation (**Fig. 5B**). It is possible that the shift of the nascent chain upon distortion of the P-site tRNA by PoxtA also contributes to “brushing” the drug from the ribosome (Ousalem et al., 2019), analogous to the mechanism proposed for leader peptides that confer resistance to macrolides (Tenson and Mankin, 2001). We also cannot exclude the possibility that conformational changes in PoxtA upon ATP hydrolysis play a role in drug release, in addition to facilitating dissociation of PoxtA from the ribosome (**Fig. 5C**).

What prevents PhO antibiotics from rebinding following their dissociation by PoxtA? As suggested previously for other ARE-ABCFs (Crowe-McAuliffe et al., 2018; Crowe-McAuliffe et al., 2021), we favour a model where, following PoxtA dissociation, the peptidyl-tRNA can reaccommodate in the P site, thus allowing accommodation of a tRNA in the A site, which, in turn, would occlude the PhO binding site (**Fig. 5D**). In this respect, we note that in contrast to previous ARE-ABCF structures (Crowe-McAuliffe et al., 2018; Crowe-McAuliffe et al., 2021), we observe here a subpopulation of PoxtA-70S complexes where the A-site tRNA is fully accommodated at the PTC despite the presence of PoxtA and the distorted P-site tRNA. Thus, once the drug has been released, the A-site tRNA could accommodate at the PTC and following peptide bond formation and translocation, the context of the nascent chain would be shifted by one amino acid i.e. alanine in the −1 position would now be located at the −2 position and therefore disfavour drug rebinding (**Fig. 5D**). PoxtA and OptrA are often found on the same mobile genetic element as drug efflux pumps (Schwarz et al., 2021), which may also contribute to preventing drug rebinding by transporting the dissociated drug directly out of the cell.

Our insights into the mechanism of action of PoxtA also provide a possible explanation for why these ARE-ABCFs do not confer resistance to MS_B_ or PLS_A_ antibiotics. Firstly, the ARD of PoxtA is too short to sterically overlap with these drugs and directly displace them from the ribosome, and secondly, binding of PoxtA does not perturb the rRNA portion of the drug binding sites and therefore could not induce their dissociation by conformational relays in the 23S rRNA. For PLS_A_ antibiotics, which stall translation after initiation, there is no nascent chain that forms part of the drug-binding site. In the case of MS_B_ antibiotics, which stall translation elongation but bind deeper in the nascent peptide exit channel, perhaps the “pulling” effect of PoxtA on the nascent chain is mitigated by conformation elasticity of the several amino acids between the P-tRNA and the drug-binding site. Interestingly, ARE-ABCF antibiotics, such as MsrE or LsaA that confer resistance to MS_B_ and PLS_A_ antibiotics, respectively, do not confer resistance to PhO antibiotics, despite direct overlap between the ARDs and the drug binding sites. We speculate that in these cases, the peptidyl-tRNA of the PhO-stalled ribosomes are refractory to the action of these ARE-ABCFs to distort the P-site tRNA; however, further investigations will be needed to validate this.

Given the similarity in ARD length and antibiotic spectrum, we believe that the findings and model presented here for PoxtA are likely to also be applicable for OptrA. Indeed, like OptrA, our MIC analysis provides no evidence for PoxtA conferring resistance to tetracycline antibiotics, consistent with the binding site of PoxtA in the E site being far from the tetracycline binding site located in the decoding A-site of the small subunit. We also see no evidence for conformational differences in the 16S rRNA nucleotides that comprise the tetracycline binding site in the absence or presence of PoxtA. For this reason, we suggest reassigning letters from the PoxtA acronym from <u>p</u>henicol <u>ox</u>azolidinone <u>t</u>etracycline_A to <u>p</u>henicol <u>ox</u>azolidinone <u>t</u>ransmissible A, analogous to OptrA.

## Methods

### *Identification of* poxtA *EF9F6 and characterisation of* E. faecium *9-F-6 antibiotic susceptibility*

The *poxtA* EF9F6 gene was identified in the multidrug-resistant ST872 *E. faecium* 9-F-6 isolated in 2012 from faeces of a patient in a Norwegian hospital which had recently also been hospitalized in India (Sivertsen et al., 2018). ST872 is a single locus variant of ST80 which is a pandemic hospital-adapted genetic lineage. Transferable linezolid resistance (*optrA* and *cfr(D)* has recently been described in blood culture *E. faecium* ST872 strain from Australia (Pang et al., 2020). Species identification was performed by MALDI-TOF (Bruker, Billerica, US) according to the manufacturer’s instructions and later confirmed by whole genome sequencing (Sivertsen et al., 2018). Antimicrobial susceptibility testing was performed by broth microdilution using the EUENCF Sensititre plate (Thermo Fisher Scientific, Waltham, Massachusetts, USA), and further by gradient tests for vancomycin, teicoplanin, ampicillin, clindamycin, chloramphenicol and gentamicin (MIC Test strip, Liofilchem, Roseto Degli Abruzzi, Italy). The results (MICs) were interpreted according to EUCAST clinical breakpoints v. 10.0 2020 and MICs were within the accepted range for quality control strain *E. faecalis* ATCC 29212. The *E. faecium* 9-F-6 is resistant to linezolid (MIC = 8 mg/L), ampicillin (>256 mg/L), ciprofloxacin (> 16 mg/L), high-level gentamicin (HLGR >256 mg/L), high-level streptomycin (HLSR >1024 mg/L) and high-level glycopeptides (vancomycin >256 mg/L and teicoplanin 128 mg/L), but susceptible to quinupristin/dalfopristin (MIC 4 mg/L) and tigecycline (MIC 0,12 mg/L) (**Table S1**). Whole-genome sequences confirmed *vanA* and the *aac(6’)*-*Ie-aph(2’’)-Ia* genes conferring high-level glycopeptide and gentamicin resistance, respectively, as well as a defunct, functionally inactive *cfr* pseudogene.

### Strains and plasmids

All bacterial strains and plasmids used in the study are listed in **Table S3**. *E. faecalis* TX5332 (Rif^r^ Fus^r^ Kan^r^; *lsa* gene disruption mutant (OG1RF *lsa*::pTEX4577)) is a *lsa* knockout strain that was received from Barbara Murray (Singh et al., 2002). pCIE_spec_ plasmid was constructed on the basis of the pCIE_cam_ plasmid, where the chloramphenicol resistance gene was substituted with spectinomycin, due to intrinsic resistance of PoxtA to phenicol antibiotics. All cloning was performed by the PEP facility (Umeå University). Plasmids encoding OptrA-ST16 (Vorobieva et al., 2017) and OptrA-E35048 (Morroni et al., 2018) were kindly provided by Anette M. Hammerum and Alberto Antonelli, respectively. In each case, the *optrA* gene was amplified with ribosome binding sequence (RBS) (TAAGAGGAGGAGATAAAC) and inserted into pCIE_spec_ plasmid, resulting in pCIE_spec_:*optrA-ST16* and pCIE_spec_:*optrA-E35048*, respectively. The *poxtA AOUC-0915* gene (Antonelli et al., 2018) was PCR amplified from genomic DNA of *S. aureus* AOUC 0915 (kindly provided by Alberto Antonelli) and inserted in pCIE_spec_ plasmid using the BamHI and HindIII restriction sites. The RBS (TAAGAGGAGGAGATAAAC) was inserted directly in front of the ATG to improve expression levels. Genetic material encoding PoxtA-E9F6 (OZN12776.1) was obtained from an *E. faecium* (9-F-6) isolated from a Norwegian patient in 2012, the genome of which has been sequenced (GCA_002263195.1) (Sivertsen et al., 2018). The *poxtA-E9F6* gene with added RBS was inserted in the pCIE_spec_ plasmid resulting in pCIE_spec_:*poxtA-EF9F6*. These plasmids were used for MIC determinations. For use in pull-out experiments we introduced His_6_-TEV-FLAG_3_ tags at the C-terminus of either OptrA or PoxtA protein, with and without EQ mutations (E184Q and E471Q for PoxtA EQ_2_, E190Q and E470Q for OptrA EQ_2_, E470Q for OptrA EQ_1_) expressed from pCIE_cam_ plasmid.

### Bacterial transformation

*E. faecalis* electrocompetent cells were prepared as described previously (Bhardwaj et al., 2016). Briefly, an over-night culture *E. faecalis* TX5332 grown in the presence of 2 mg/mL of kanamycin was diluted to OD_600_ of 0.05 in 50 mL of Brain Heart Infusion (BHI) media and grown further to an OD_600_ of 0.6-0.7 at 37 °C with moderate shaking (160 rpm). Cells were collected by centrifugation at 4,000 rpm at 4 °C for 10 min. Cells were resuspended in 0.5 mL of sterile lysozyme buffer (10 mM Tris-HCl pH 8; 50 mM NaCl, 10 mM EDTA, 35 µg/mL lysozyme), transferred to 0.5 mL Eppendorf tube and incubated at 37 °C for 30 minutes. Cells were pelleted at 10000 rpm at 4 °C for 10 min and washed three times with 1.5 mL of ice-cold electroporation buffer (0.5 M sucrose, 10% (w/v) glycerol). After last wash the cells were resuspended in 500 µL of ice-cold electroporation buffer and aliquoted and stored at – 80°C. For electroporation, 35 µL of electrocompetent cells were supplemented with 0.5–1 µg of plasmid DNA, transferred to ice-cold 1 mm electroporation cuvette and electroporated at 1.8 keV. Immediately after electroporation 1 mL of ice-cold BHI media was added to the cells, the content of the cuvette was transferred to 1.5 mL Eppendorf tubes and the cells were recovered at 37 °C for 2.5 hours and plated to BHI plates containing appropriate antibiotics.

### Antibiotic susceptibility testing

*E. faecalis* cells were grown in BHI media supplemented with 2 mg/mL kanamycin (to prevent *lsa* revertants), 0.1 mg/mL spectinomycin (to maintain the pCIE_spec_ plasmid), 100 ng/mL of cCF10 peptide (to induce expression of proteins of interest) as well as increasing concentrations of antibiotics. The media was inoculated with 5 × 10^5^ CFU/mL (OD_600_ of approximately 0.0005) of *E. faecalis* Δ*lsaA* (*lsa::Kan*) strain TX5332 transformed either with empty pCIE_spec_ plasmid or with pCIE_spec_ encoding indicated protein of interest. After 16-20 h at 37 °C without shaking, the presence or absence of bacterial growth was scored by eye.

### Preparation of bacterial biomass

*E. faecalis* TX5332 transformed with pCIE_spec_ / pCIE_cam_-based expression constructs (either empty vector or expressing wild type/EQ_2_ variants of both PoxtA-HTF and OptrA-HTF as well as EQ_1_ variant of OptrA-HTF) were grown overnight from single colony in BHI media supplemented with appropriate antibiotics (100 µg/mL of spectinomycin for pCIE_spec_-based constructs, 10 µg/mL chloramphenicol for pCIE_cam_-based constructs). Overnight cultures were then diluted to a starting OD_600_ of 0.05 in 200 mL of the same media. Cells were grown with intensive shaking at 37 °C till OD_600_ of 0.6 and were induced with 100 ng/mL of cCF10 peptide for 30 minutes prior harvesting by centrifugation at 10,000 x g for 15 min at 4 °C.

### Preparation of clarified lysates

Cell pellets were resuspended in 1.5 mL of cell opening buffer (95 mM KCl, 5 mM NH_4_Cl, 20 mM HEPES pH 7.5, 1 mM DTT, 5 mM Mg(OAc)_2_, 0.5 mM CaCl_2_, 8 mM putrescine, 0.5-0.75 mM ATP, 1 mM spermidine, 1 tablet of cOmplete™ EDTA-free Protease Inhibitor Cocktail (Roche) per 10 mL of buffer). Resuspended cells were opened by FastPrep homogeniser (MP Biomedicals) with 0.1 mm Zirconium beads (Techtum) in 4 cycles by 20 seconds with a 1 min chill on ice. Cell debris was removed after centrifugation at 14,800 x g for 15 min at 4 °C. Total protein concentration in supernatant was measured by Bradford assay (BioRad), supernatant was aliquoted and frozen in liquid nitrogen.

### Affinity purification on anti-FLAG M2 affinity gel

100 µL of well-mixed anti-FLAG M2 Affinity Gel aliquots (Sigma) were loaded on columns (Micro Bio-Spin Columns, Bio-Rad) and washed two times with 1 mL of cell opening buffer by gravity flow. All incubations, washings and elutions were done at 4 °C. The total protein concentration of each lysate was adjusted to 2 mg/mL with cell opening buffer with HEPES pH 7.5 to glycine pH 9.0. 1 mL of each lysate was loaded on columns and incubated for 2 h with end-over-end mixing for binding. The columns were washed 5 times with 1 mL of cell opening buffer by gravity flow. For elution of FLAG-tagged proteins and their complexes 100 µL of 0.2 mg/mL FLAG_3_ peptide (Sigma) was added to samples, the solutions were incubated at 4 °C for 20 minutes with end-over-end mixing. Elutions were collected by centrifugation at 2,000 x g for 2 min at 4 °C. 20 µL aliquots of collected samples (flow-through, washes and elutions) were mixed with 5 µL of 5 x SDS loading buffer and heated up at 95 °C for 15 min. The beads remaining in the column were resuspended in 100 µL of 1x SDS loading buffer. Denatured samples were loaded on 15% SDS-PAGE. SDS-gels were stained by “Blue-Silver” Coomassie Staining (Candiano et al., 2004) and destained in water overnight hours or overnight before imaging with LAS4000 (GE Healthcare).

### Preparation of cryo-EM grids

Elutions from pull-downs were kept on ice until being applied within two hours to glow discharged cryo-grids (Quantifoil 2/2 Cu_300_ coated with 2 nm continuous carbon). 3.5 µL of sample was loaded on grids 3 times manually at room temperature conditions and a fourth time in Vitrobot (FEI) under conditions of 100% humidity at 4 °C, blotted for 5 seconds and vitrified by plunge-freezing in liquid ethane. Samples were imaged on a Titan Krios (FEI) operated at 300 kV at a nominal magnification of 165,000 x (0.86 Å/pixel) with a Gatan K2 Summit camera at an exposure rate of 5.85 electrons/pixel/s with a 4 seconds exposure and 20 frames using the EPU software.

### Single-particle reconstruction

Processing was performed in RELION 3.1 unless otherwise specified (Zivanov et al., 2018). MotionCor2 with 5×5 patches and CTFFIND4 (using power spectra) were used for motion correction and initial CTF estimation (Rohou and Grigorieff, 2015; Zheng et al., 2017). Micrographs with estimated CTF fits beyond 4.5 Å and CTF figure of merits > 0.04 were selected for further processing. Particles were picked with crYOLO using the general model (Wagner et al., 2019). After 2D classification, all ribosome-like classes were selected, particles extracted with a 3× reduced pixel size (2.46 Å), and an initial model created *ab initio*. After 3D refinement using the *ab initio* model as a reference (Scheres and Chen, 2012), 3D classification with eight classes and without angular searching was performed. The majority of particles (∼80%) clustered into two classes that contained protein-like density in the E site, and which were selected for further processing. Particles were re-extracted at the original pixel size, and serial 3D refinements with CTF refinement and Bayesian polishing (Zivanov et al., 2019) were performed until the resolution did not improve further, resulting in the ‘combined 70S volume’. A mask around the A, P, and E sites was created and used for partial signal subtraction with re-extraction at a pixel size of 2.46 Å. These particles were used for 3D classification with six classes, T = 40 and the resolution of the expectation step limited to 12 Å. Four of the six resulting classes, labelled states I–IV, were chosen for refinement with the original pixel size. For the multibody refinement of the combined 70S class (Nakane et al., 2018), volumes corresponding to the LSU core, CP, SSU body, SSU head, and E-site were isolated using the volume eraser tool in UCSF ChimeraX (Pettersen et al., 2021), and masks created from the densities low-pass-filtered to 30 Å. Bsoft was used to estimate local resolution (Heymann, 2018).

### Molecular modelling

Molecular models were created/adjusted with Coot (Casanal et al., 2020) and ISOLDE (Croll, 2018), and refined with Phenix (Liebschner et al., 2019) against unsharpened maps. A previous structure of the *E. faecalis* 70S ribosome in complex with the ABCF-ARE LsaA (PDB 7NHK) (Crowe-McAuliffe et al., 2021) was used as the starting model for the ribosome, initially into multibody-refined maps from the combined 70S volume. PDB IDs 7K00 (Watson et al., 2020) and 6O90 (Murphy et al., 2020) were also used as templates in parts of the 70S, and PDB ID 3U4M (Tishchenko et al., 2012) was used as a template for the L1 stalk region. Likely metal ions were assigned by the presence of typical coordination shapes, as well as strength of density and estimated coordination distances. However, we caution that these assignments could not always be made unambiguously. Phenix.douse was used to place water molecules. A recent structure of the *S. aureus* ribosome with modified nucleotides (PDB ID 6YEF (Golubev et al., 2020)) was compared to the density to check for RNA modifications, and where the density matched the modification with high confidence, that modification was inserted into the *E. faecalis* 70S model. For PoxtA, homology models were generated by SWISS-MODEL (Waterhouse et al., 2018) using the crystal structure of EttA (PDB ID 4FIN (Boel et al., 2014)) and the previous cryo-EM structure of LsaA (PDB 7NHK) (Crowe-McAuliffe et al., 2021). The ARD was sufficiently well-resolved that it could be built manually. The Arm insertion in NBD1 was built manually with low confidence (especially in the loop connecting the two α-helices). PSI-PRED secondary structure predictions were used to help define secondary structure boundaries (Buchan and Jones, 2019). The models for states I–IV were assembled using the model from the combined 70S as a template. For the A-tRNA in state III, tRNA-Lys from PDB ID 5E7K (Rozov et al., 2016) was used as a template for *E. faecalis* tRNA-Lys-UUU-1-1 (gtRNAdb nomenclature (Chan and Lowe, 2016)) based on density.

### Sequence analysis methods

Representative ABCF sequences were aligned with Mafft L-INS-i 7.453 (Katoh and Standley, 2013) and visualised with Jalview 2.11.1.4 (Waterhouse et al., 2009) and Aliview 1.26 (Larsson, 2014). Phylogenetic analysis was carried out with IQTree version 2.1.2 on the CIPRES server with 1000 rapid bootstrap replicates and automatic model detection (Miller et al., 2010; Minh et al., 2020). Positions with more than 50% gaps were removed with TrimAL v1.4.rev6 before phylogenetic analysis (Capella-Gutierrez et al., 2009).

### Figures

Figures were created using UCSF ChimeraX, PyMol v2.4, and assembled with Adobe Illustrator (Adobe Inc.).

## Data availability

Micrographs have been deposited as uncorrected frames in the Electron Microscopy Public Image Archive (EMPIAR) with the accession codes EMPIAR-10XXX [https://www.ebi.ac.uk/pdbe/emdb/empiar/entry/10XXX/]. Cryo-EM maps have been deposited in the Electron Microscopy Data Bank (EMDB) with accession codes EMD-12XXX [https://www.ebi.ac.uk/pdbe/entry/emdb/EMD-12XXX] (PoxtA-70S state I), EMD-12XXX [https://www.ebi.ac.uk/pdbe/entry/emdb/EMD-12XXX] (PoxtA-70S state II), EMD-12XXX [https://www.ebi.ac.uk/pdbe/entry/emdb/EMD-12XXX] (PoxtA-70S state III with A-site tRNA) and EMD-12XXX [https://www.ebi.ac.uk/pdbe/entry/emdb/EMD-12XXX] (*E. faecalis* 70S with P-tRNA state IV). Molecular models have been deposited in the Protein Data Bank with accession codes 7XXX [https://doi.org/10.2210/pdb7XXX/pdb] (PoxtA-70S state I), 7XXX [https://doi.org/10.2210/pdb7XXX/pdb] (PoxtA-70S state II), 7XXX [https://doi.org/10.2210/pdb7XXX/pdb] (PoxtA-70S state III with A-site tRNA) and 7XXX [https://doi.org/10.2210/pdb7XXX/pdb] (*E. faecalis* 70S with P-tRNA, state IV). Source data are provided with this paper.

## Acknowledgments

We are grateful to Barbara E. Murray for sharing *E. faecalis* Δ*lsaA* (*lsa*::Kan) strain TX5332 (Singh et al., 2002), Jose A. Lemos for sharing pCIE plasmid (Weaver et al., 2017), Alberto Antonelli for sharing genetic material containing *poxtA* AOUC-0915 and *optrA* E35048 and Anette M. Hammerum for sharing the plasmid containing *optrA* ST16 (Vorobieva et al., 2017). We thank Michael Hall for help with cryo-EM data collection. The electron microscopy data was collected at the Umeå Core Facility for Electron Microscopy, a node of the Cryo-EM Swedish National Facility, funded by the Knut and Alice Wallenberg, Family Erling Persson and Kempe Foundations, SciLifeLab, Stockholm University and Umeå University. This work was supported by the Deutsche Forschungsgemeinschaft (DFG) (grant WI3285/8-1 to D.N.W), Swedish Research Council (Vetenskapsrådet) grants (2017-03783 to V.H. and 2019-01085 to G.C.A.), Ragnar Söderbergs Stiftelse (to V.H.), postdoctoral grant from the Umeå Centre for Microbial Research, UCMR (to H.T.), the European Union from the European Regional Development Fund through the Centre of Excellence in Molecular Cell Engineering (2014-2020.4.01.15-0013 to V.H.); and the Estonian Research Council (PRG335 to V.H.). D.N.W. and V.H. groups are also supported by the Deutsche Zentrum für Luft-und Raumfahrt (DLR01Kl1820 to D.N.W.) and the Swedish Research Council (2018-00956 to V.H.) within the RIBOTARGET consortium under the framework of JPIAMR. GCA and VH were also supported by a project grant from the Knut and Alice Wallenberg Foundation (2020-0037 to GCA). Y.S.P. group is supported by the National Institutes of Health (R01-GM132302 to Y.S.P).

## Author contributions

C.C.M. processed the microscopy data, generated and refined the molecular models and made the structure figures. V.M, H.T. and M.K. cloned the ARE constructs, performed genetic manipulations of *E. faecalis*, performed polysome fractionations and immunoblotting as well as performed MICs. V.M, K.J.T. and M.K. performed immunoprecipitations. V.M. prepared cryo-EM grids and collected cryo-EM datasets. A.S. and K.H. provided research materials and performed MICs. G.C.A. performed sequence conservation analyses. Y.S.P. provided structural insights into the mechanism of chloramphenicol action. C.C.M. and D.N.W. wrote the manuscript with input from all authors. D.N.W and V.H. conceived and supervised the project.

## Supplementary Tables

**Table S1.**
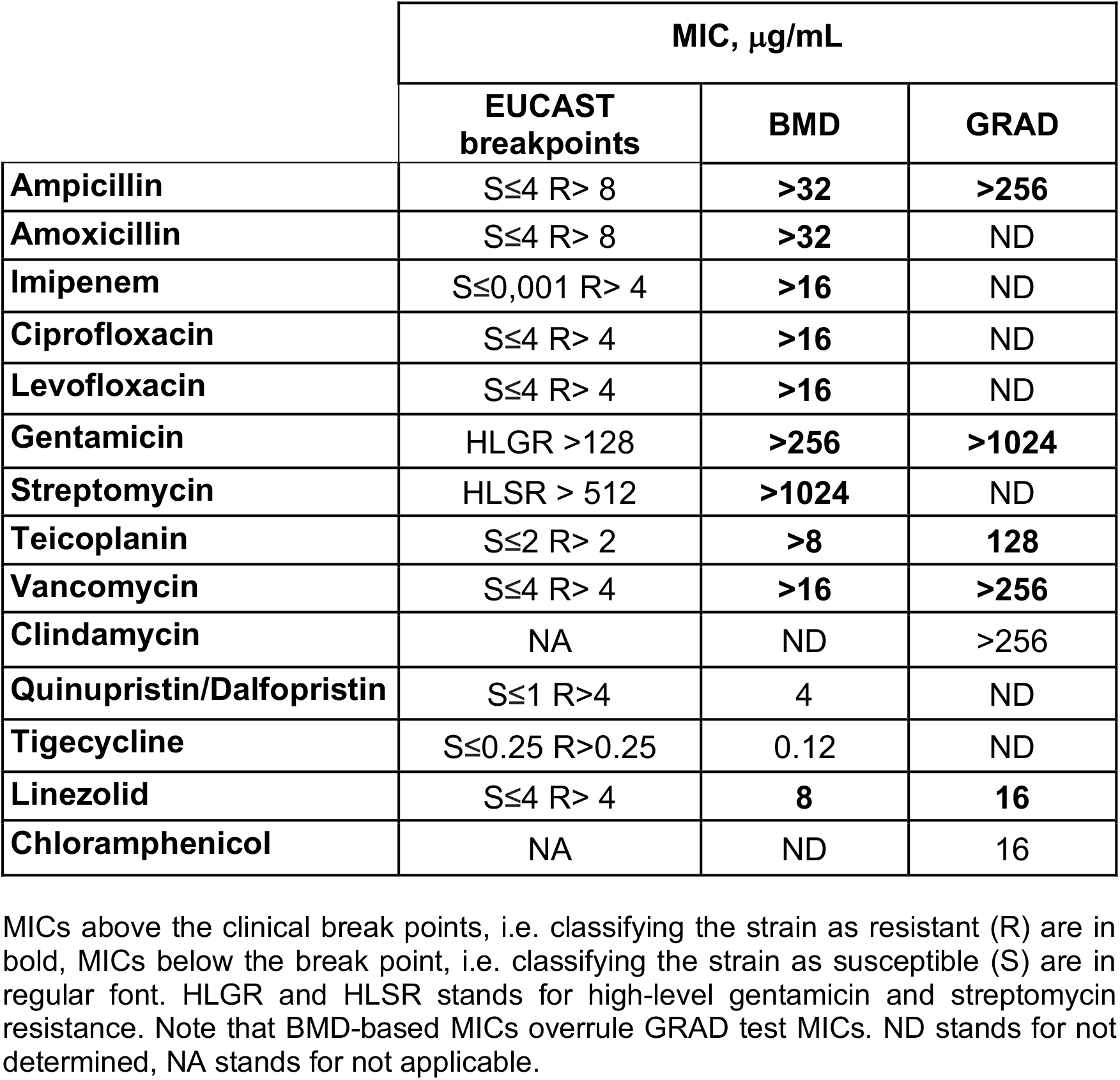
Broth microdilution (BMD) and gradient (GRAD) MIC testing of multidrug-resistant ST872 *E. faecium* 9-F-6.

**Table S2.** Cryo-EM data collection, modelling and refinement statistics.

|  | **Combined**<br>**70S** | **State I** | **State II** | **State III** | **State IV** |
| --- | --- | --- | --- | --- | --- |
| **Data collection** |  |  |  |  |  |
| Magnification (×) | 165 000 | 165 000 | 165 000 | 165 000 | 165 000 |
| Electron fluence<br>(e<sup>-</sup>/Å<sup>2</sup>) | 30.255 | 30.255 | 30.255 | 30.255 | 30.255 |
| Defocus range<br>(µm) | -0.5 to<br>-1.5 | -0.5 to<br>-1.5 | -0.5 to<br>-1.5 | -0.5 to<br>-1.5 | -0.5 to<br>-1.5 |
| Pixel size (Å) | 0.82 | 0.82 | 0.82 | 0.82 | 0.82 |
| Initial particles | 203 321 | 203 321 | 203 321 | 203 321 | 203 321 |
| Final particles | 112 877 | 29 581 | 29 875 | 23 806 | 18 512 |
| Average resolution<br>(Å) (FSC threshold<br>0.143) | 2.4 | 2.9 | 3.0 | 2.9 | 3.1 |
| **Model**<br>**composition** |  |  |  |  |  |
| Atoms | 151 224 | 147 584 | 147 584 | 149 091 | 139 841 |
| Protein residues | 6 097 | 6 097 | 6 097 | 6 097 | 5 337 |
| RNA bases | 4 625 | 4 625 | 4 625 | 4697 | 4 549 |
| **Refinement** |  |  |  |  |  |
| Map CC around<br>atoms | 0.80 | 0.71 | 0.69 | 0.73 | 0.72 |
| Map CC whole unit<br>cell | 0.80 | 0.70 | 0.69 | 0.71 | 0.70 |
| Map sharpening B<br>factor (Å<sup>2</sup>) | -46 | -55 | -59 | -58 | -59 |
| R.M.S. deviations |  |  |  |  |  |
| Bond lengths<br>(Å) | 0.007 | 0.007 | 0.004 | 0.005 | 0.004 |
| Bond angles (°) | 1.041 | 0.861 | 0.677 | 0.743 | 0.500 |
| **Validation** |  |  |  |  |  |
| MolProbity score | 1.76 | 1.95 | 1.90 | 1.81 | 1.48 |
| Clash score | 4.38 | 6.11 | 6.76 | 6.06 | 5.98 |
| Poor rotamers (%) | 2.88 | 3.58 | 3.19 | 2.80 | 1.12 |
| Ramachandran<br>statistics |  |  |  |  |  |
| Favoured (%) | 96.82 | 96.78 | 97.25 | 97.25 | 97.44 |
| Outlier (%) | 0.08 | 0.07 | 0.03 | 0.05 | 0.06 |

**Table S3.**
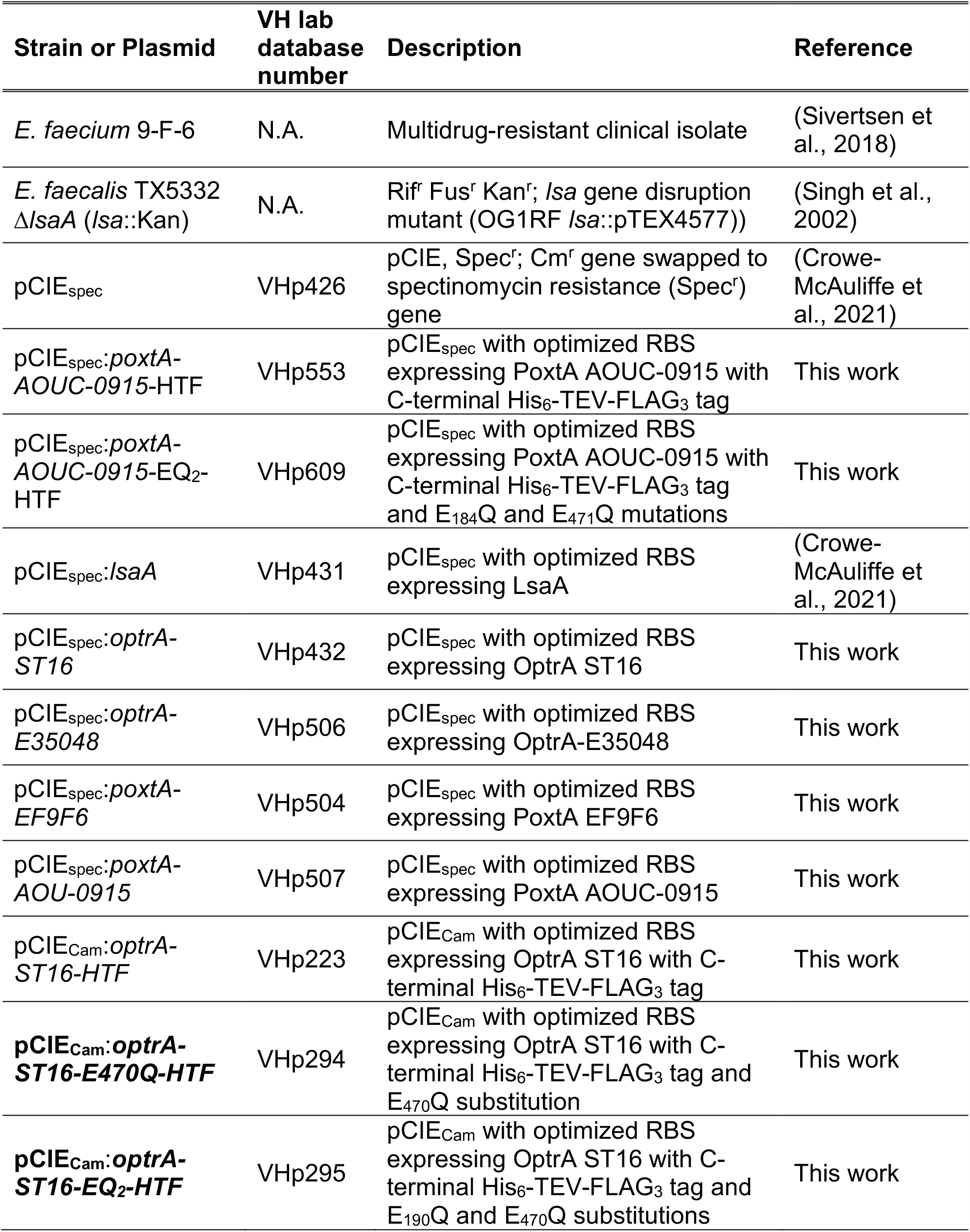
Strains and plasmids used in this study. Abbreviations used: spec – spectinomycin, RBS – ribosome binding site, cm – chloramphenicol, HTF- His_6_-TEV-FLAG_3_, ARD – antibiotic resistance domain. NA stands for not applicable.

## Supplementary Figures

**Fig. S1.**
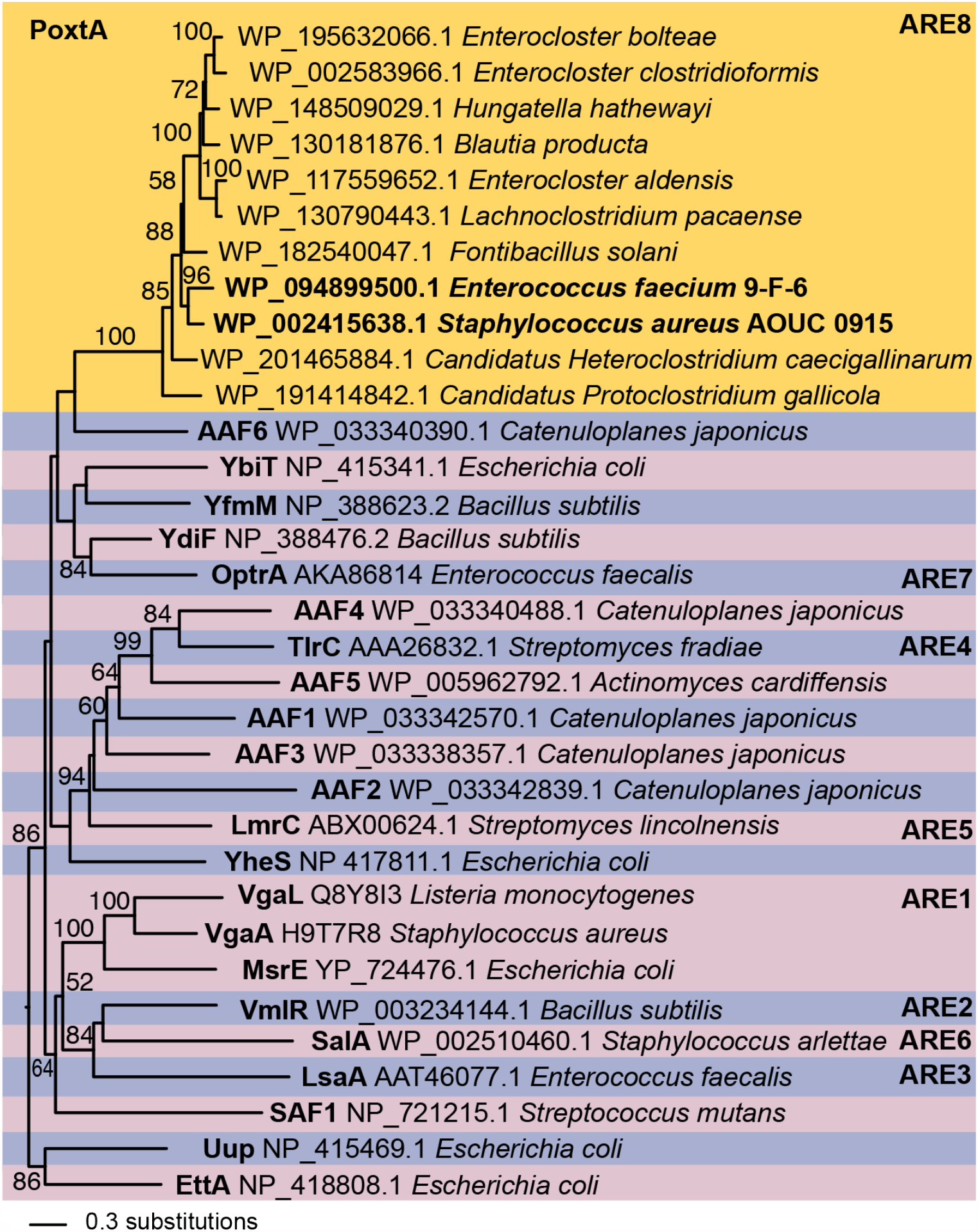
ARE-ABCF phylogeny with a focus on PoxtA. The tree is a maximum likelihood phylogeny of selected ABCF representatives. Numbers on branches correspond to IQ-Tree ultra-fast bootstrap support in percentages (Minh et al., 2020), and branch length is proportional to the number of substitutions as per the scale bar in the lower left. The ARE8 subfamily comprising PoxtA-like proteins has 100% support, but its relationship with other subfamilies is unresolved. OptrA is most closely related to the YdiF subfamily of mostly vertically inherited and presumably housekeeping ABCF (84% bootstrap support). This suggests that the similar resistance spectrum of OptrA and PoxtA is a case of convergent evolution.

**Fig. S2.**
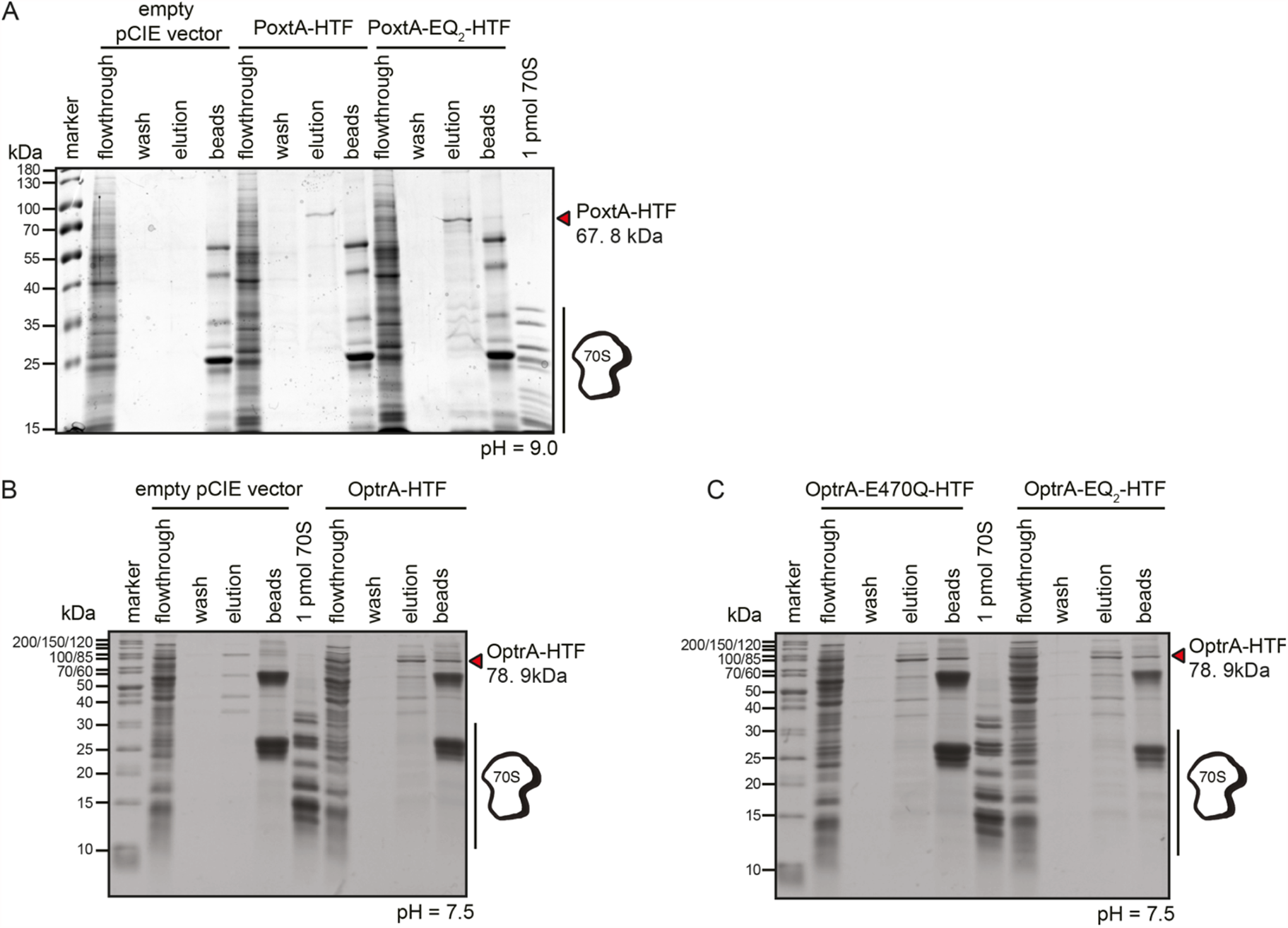
Characterization of PoxtA-E9F6 interactions with ribosomes and preparation of samples for cryo-EM reconstructions. (*A*) Affinity purification of PoxtA-EQ_2_-HTF ectopically expressed in *E. faecalis* Δ*lsaA* (*lsa*::Kan) strain TX5332. Pull-down experiments were performed in the presence of 0.5 mM ATP using clarified lysates of *E. faecalis* either transformed with empty integrative pCIE_*spec*_ vector (background control), or either expressing PoxtA-HTF or PoxtA-EQ_2_-HTF. Samples: marker: 2 μL of molecular weight marker; flowthrough: 2 μL of flowthrough; wash: 10 μL of last wash before specific elution; elution: 10 μL of elution with FLAG_3_ peptide at pH 9.0; beads: 2 μL of SDS-treated post-elution anti-FLAG beads; 70S: purified *E. faecalis* 70S ribosomes, the samples were resolved on 12 % SDS-PAGE gel. (*B, C*) Affinity purification attempts with wild-type, EQ_2_ and EQ_1_ (E470Q) *E. faecalis* OptrA-ST16-HTF ectopically expressed in TX5332 *E. faecalis*. Pull-down experiments were performed in the presence of 0.75 mM ATP at pH 7.5 using clarified lysates of *E. faecalis* either transformed with *E. faecalis* OptrA-ST16-HTF (VHp223) or expressing either *E. faecalis* OptrA-ST16-E470Q-HTF (VHp294) or OptrA-ST16-EQ_2_-HTF (VHp295). Samples: marker: molecular weight marker; lysate: 2 μL of clarified lysate, flowthrough: 2 μL of flow-through; wash: 10 μL of last wash before specific elution; elution: 10 μL of the elution with FLAG_3_ peptide; B: 10 μL of SDS-treated post-elution anti-FLAG beads; 70S: purified *E. faecalis* 70S ribosomes. The samples were resolved on 15% SDS-PAGE gel.

**Fig. S3.**
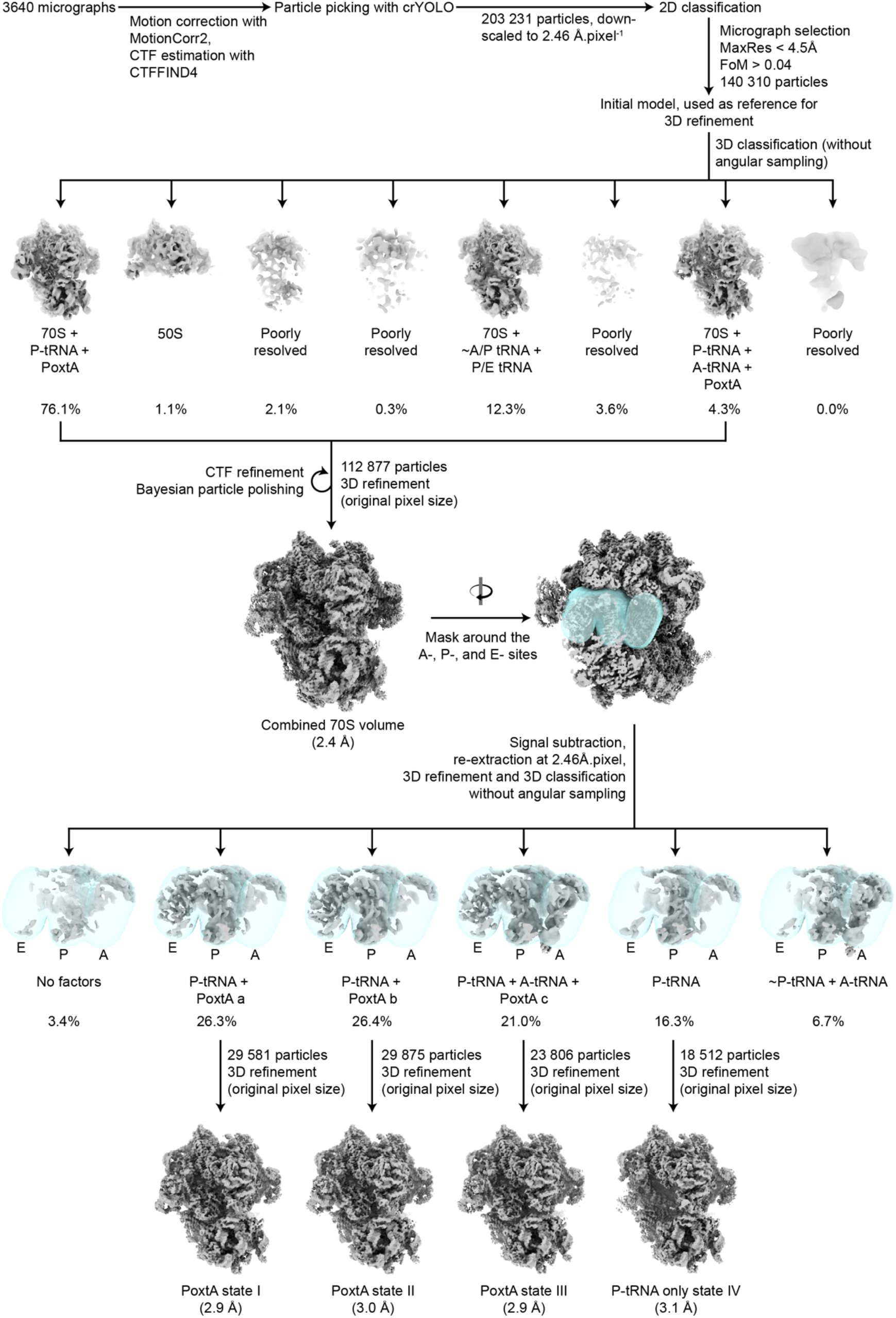
Processing scheme for the PoxtA-70S complex. See also Methods for details.

**Fig. S4.**
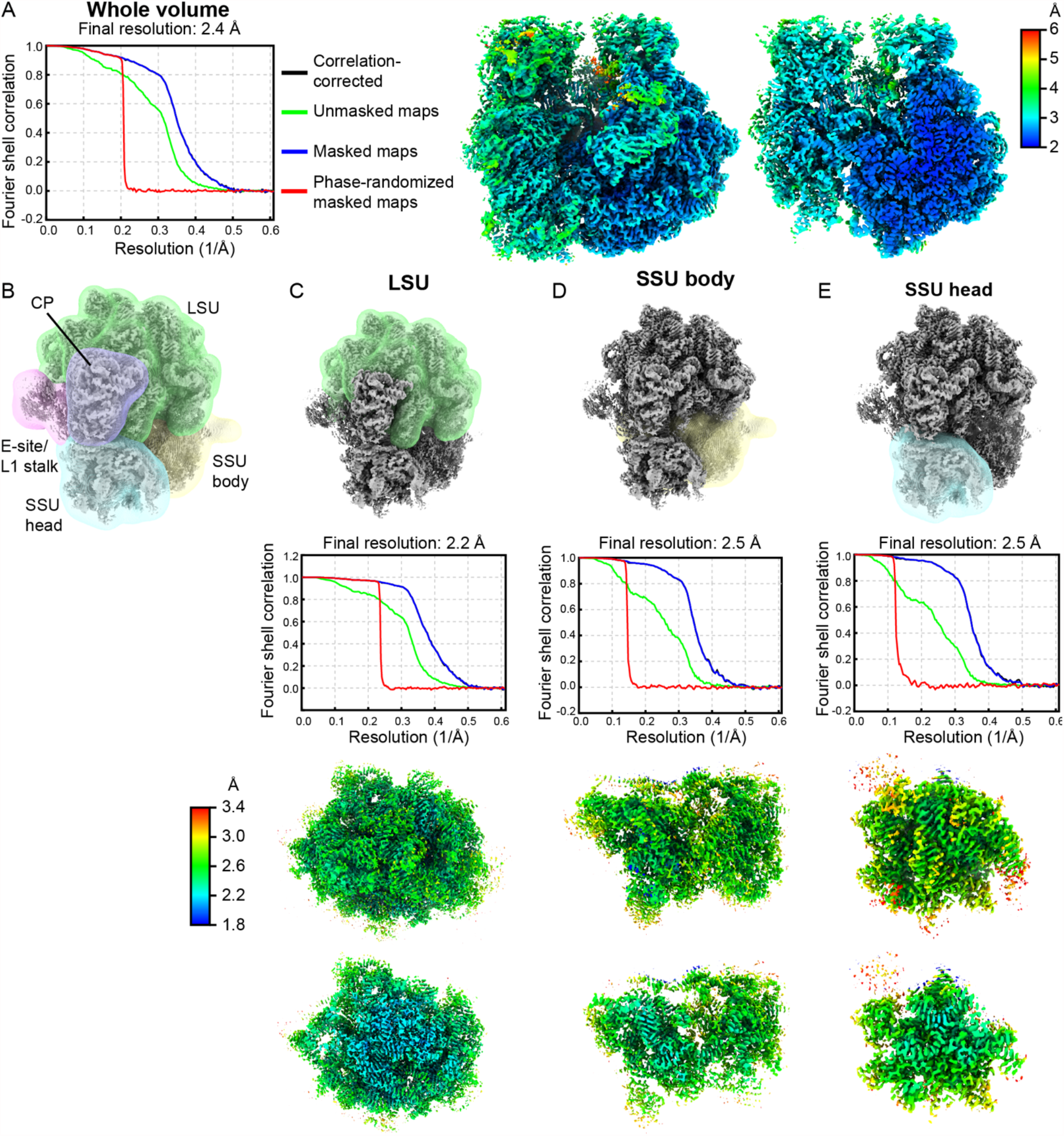
Local resolutions and multibody refinements for the combined 70S volume. (*A*) FSC curve and local resolution images for the combined 70S volume. (*B*) Overview of masks used for multibody refinement (*C*–*E*) masks relative to whole 70S (top row), FSC curves (second row), and density coloured according to local resolution (bottom two rows) for (*C*) the LSU core, (*D*) the SSU body, and (*E*) the SSU head.

**Fig. S5.**
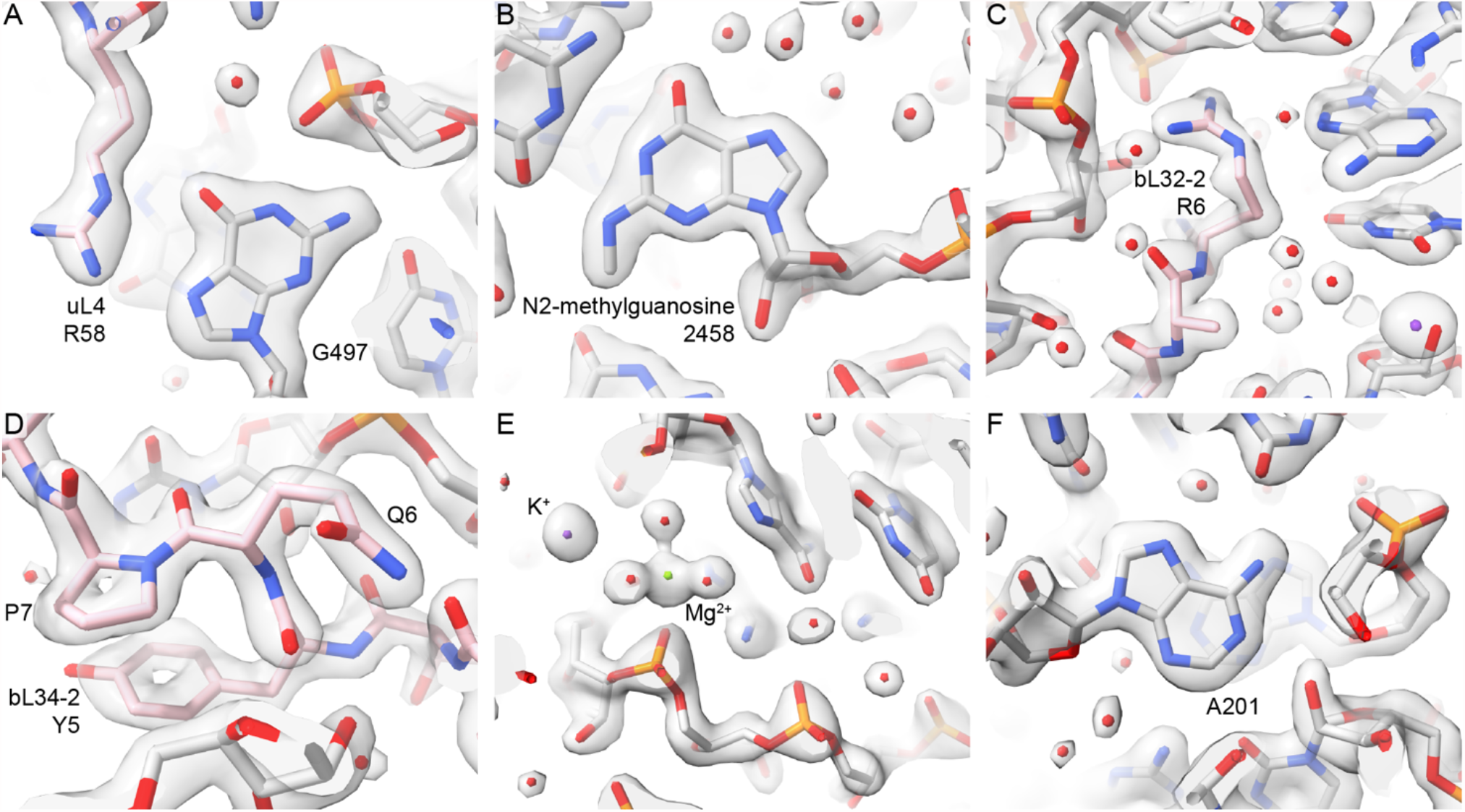
Selected density images from the high-resolution LSU core. Modelled water molecules are colored red, magnesium green, and potassium purple.

**Fig. S6.**
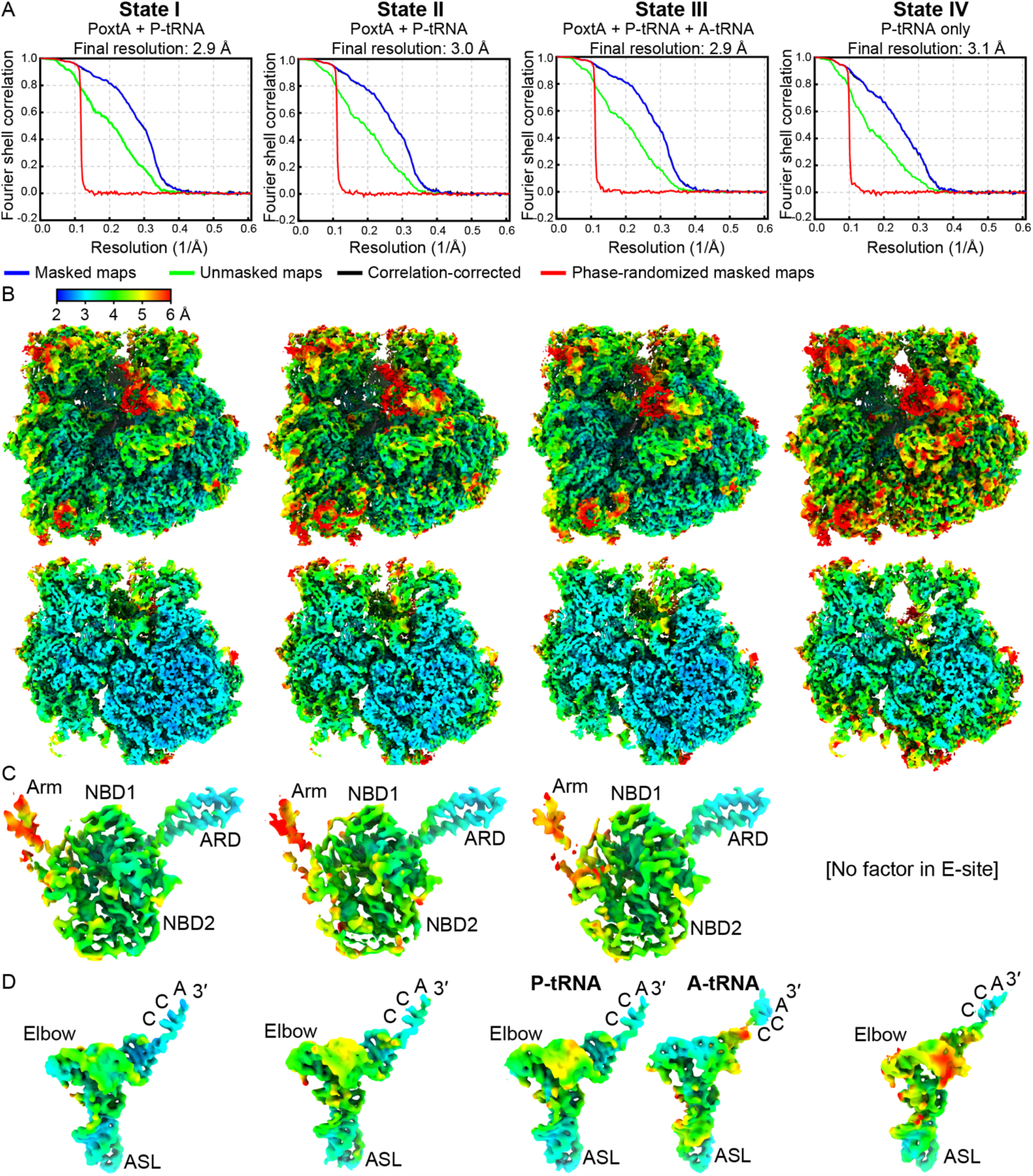
FSC curves and density coloured by local resolution for states I-IV. (*A*) FSC curves. (*B*) density coloured by local resolution for whole volumes with cut-throughs (bottom row). (*C*) Isolated density for isolated PoxtA coloured by local resolution. (*D*) Isolated density for isolated tRNAs coloured by local resolution.

**Fig. S7.**
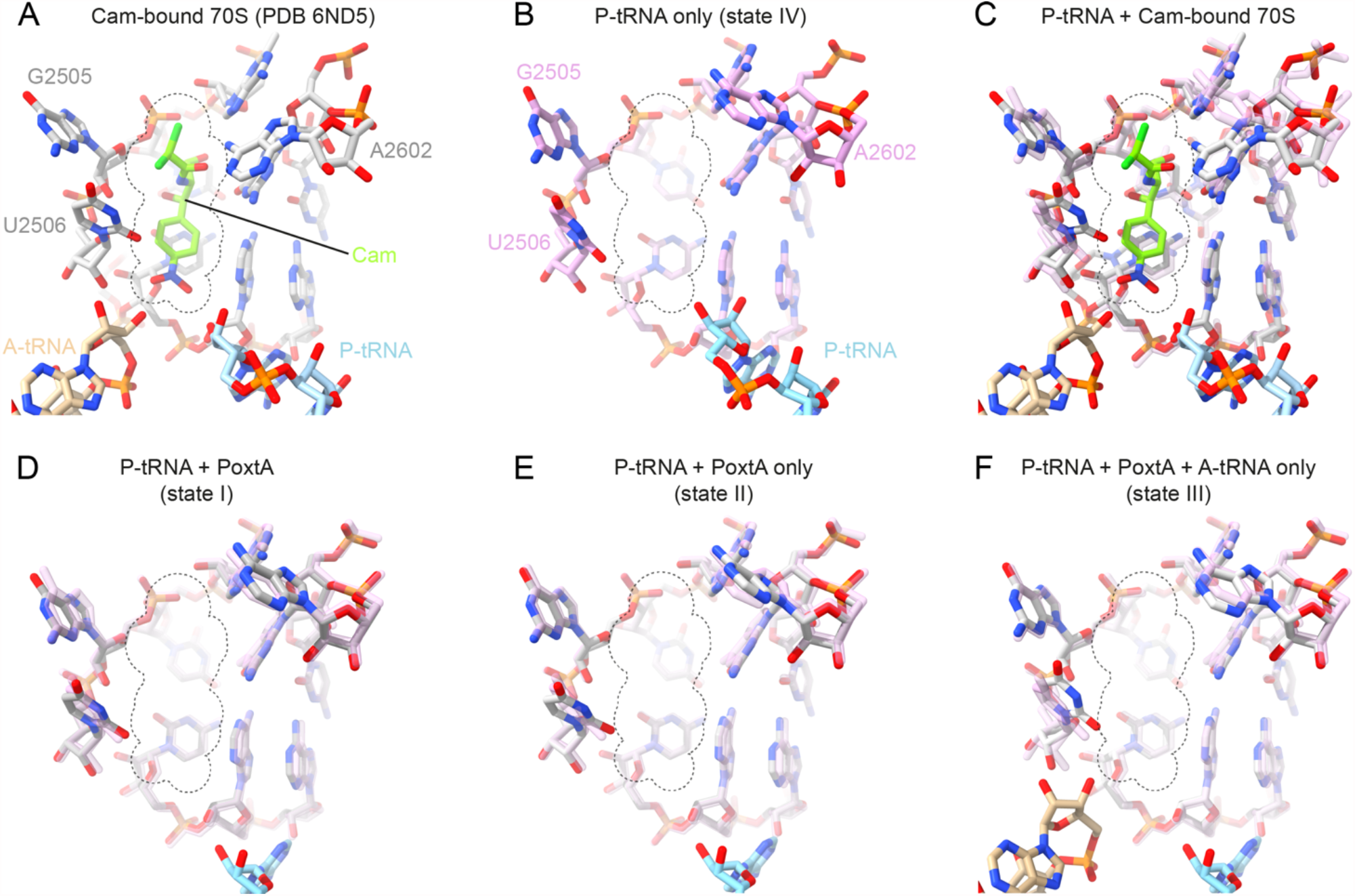
Effects of PoxtA binding on the 23S rRNA chloramphenicol binding site. (*A*) Chloramphenicol (cam, green, PDB ID 6ND5, (Svetlov et al., 2019)) bound to 23S rRNA (grey). Dotted lines indicate the extent of a space-filling model of chloramphenicol. A-tRNA (tan) and P-tRNA (light blue) are also shown). (*B*) Same view as (A) but for the P-tRNA-only bound volume (state IV). 23S rRNA is shown in pink. (*C*) Same as panel A but with P-tRNA-only (state IV) 23S rRNA superimposed (transparent pink). (*D*–*F*) As for *C*, but comparing the P-tRNA-only (state IV) 23S rRNA with PoxtA bound states I–III. The chloramphenicol outline is shown for reference.

**Fig. S8.**
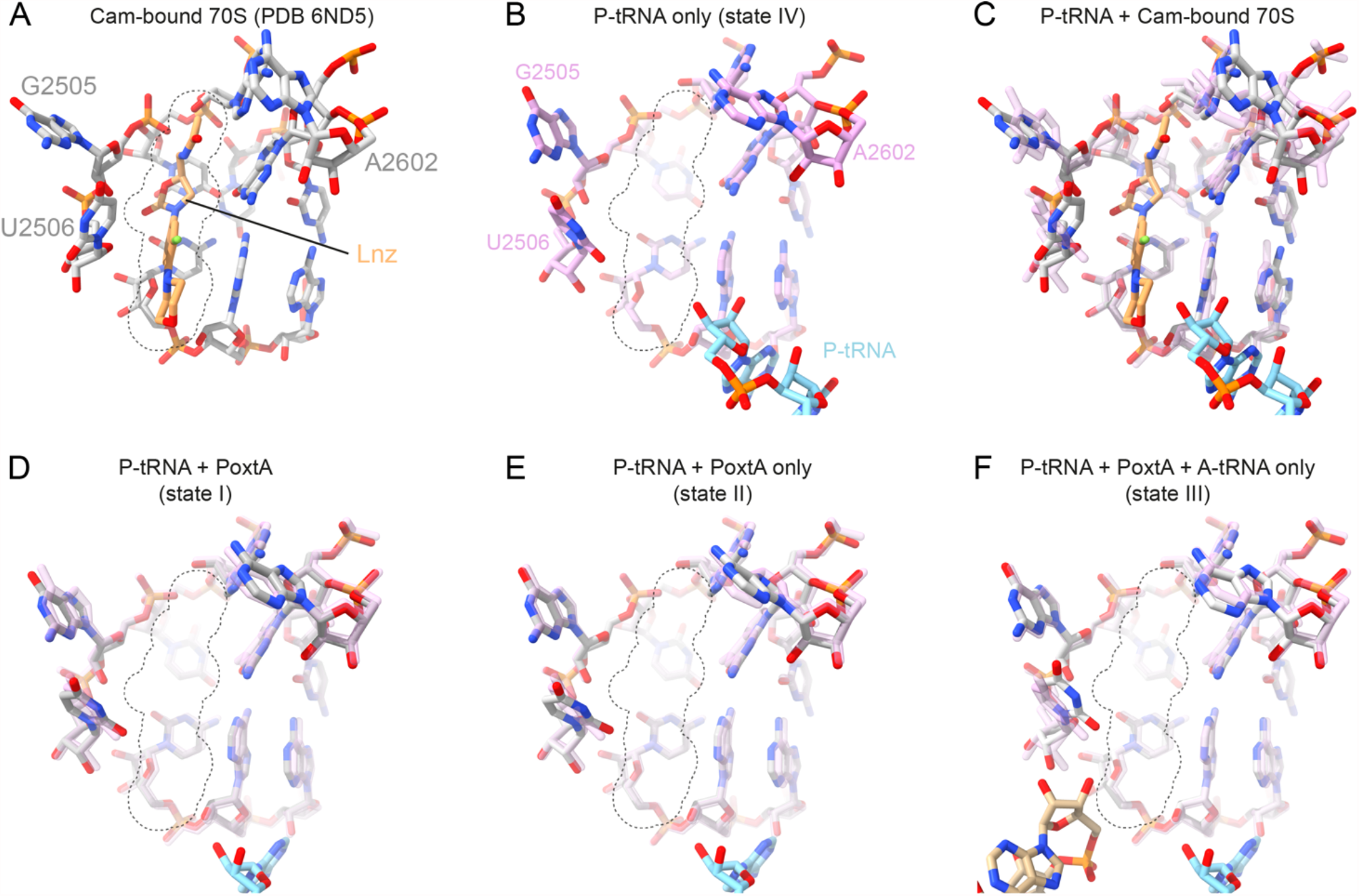
Effects of PoxtA binding on the 23S rRNA linezolid binding site. (*A*) Linezolid (Lnz, light orange, PDB ID 3DLL, (Wilson et al., 2008)) bound to 23S rRNA (grey). Dotted lines indicate the extent of a space-filling model of linezolid. A-tRNA (tan) and P-tRNA (light blue) are also shown). (*B*) Same view as (*A*) but for the P-tRNA-only bound volume (state IV). 23S rRNA is shown in pink. (*C*) Same as panel A but with P-tRNA-only (state IV) 23S rRNA superimposed (transparent pink). (*D*–*F*) As for *C*, but comparing the P-tRNA-only (state IV) 23S rRNA with PoxtA bound states I*–*III. The linezolid outline is shown for reference.

## References

Antonelli, A., D’Andrea, M.M., Brenciani, A., Galeotti, C.L., Morroni, G., Pollini, S., Varaldo, P.E., and Rossolini, G.M. (2018). Characterization of poxtA, a novel phenicol-oxazolidinone- tetracycline resistance gene from an MRSA of clinical origin. J Antimicrob Chemother 73, 1763–1769.

Bhardwaj, P., Ziegler, E., and Palmer, K.L. (2016). Chlorhexidine induces VanA-type vancomycin resistance genes in enterococci. Antimicrobial agents and chemotherapy 60, 2209–2221.

Boel, G., Smith, P.C., Ning, W., Englander, M.T., Chen, B., Hashem, Y., Testa, A.J., Fischer, J.J., Wieden, H.J., Frank, J., et al. (2014). The ABC-F protein EttA gates ribosome entry into the translation elongation cycle. Nature structural & molecular biology 21, 143–151.

Buchan, D.W.A., and Jones, D.T. (2019). The PSIPRED Protein Analysis Workbench: 20 years on. Nucleic Acids Res. 47, W402–W407.

Cai, J., Wang, Y., Schwarz, S., Lv, H., Li, Y., Liao, K., Yu, S., Zhao, K., Gu, D., Wang, X., et al. (2015). Enterococcal isolates carrying the novel oxazolidinone resistance gene optrA from hospitals in Zhejiang, Guangdong, and Henan, China, 2010-2014. Clin Microbiol Infect 21, 1095 e1091–1094.

Candiano, G., Bruschi, M., Musante, L., Santucci, L., Ghiggeri, G.M., Carnemolla, B., Orecchia, P., Zardi, L., and Righetti, P.G. (2004). Blue silver: a very sensitive colloidal Coomassie G-250 staining for proteome analysis. Electrophoresis 25, 1327–1333.

Capella-Gutierrez, S., Silla-Martinez, J.M., and Gabaldon, T. (2009). trimAl: a tool for automated alignment trimming in large-scale phylogenetic analyses. Bioinformatics 25, 1972–1973.

Casanal, A., Lohkamp, B., and Emsley, P. (2020). Current developments in Coot for macromolecular model building of Electron Cryo-microscopy and Crystallographic Data. Protein Sci 29, 1069–1078.

Chan, P.P., and Lowe, T.M. (2016). GtRNAdb 2.0: an expanded database of transfer RNA genes identified in complete and draft genomes. Nucleic Acids Res. 44, D184–189.

Chen, B., Boel, G., Hashem, Y., Ning, W., Fei, J., Wang, C., Gonzalez, R.L., Jr., Hunt, J.F., and Frank, J. (2014). EttA regulates translation by binding the ribosomal E site and restricting ribosome-tRNA dynamics. Nature structural & molecular biology 21, 152–159.

Choi, J., Marks, J., Zhang, J., Chen, D.H., Wang, J., Vazquez-Laslop, N., Mankin, A.S., and Puglisi, J.D. (2020). Dynamics of the context-specific translation arrest by chloramphenicol and linezolid. Nat Chem Biol 16, 310–317.

Croll, T.I. (2018). ISOLDE: a physically realistic environment for model building into low-resolution electron-density maps. Acta Crystallogr D Struct Biol 74, 519–530.

Crowe-McAuliffe, C., Graf, M., Huter, P., Takada, H., Abdelshahid, M., Novacek, J., Murina, V., Atkinson, G.C., Hauryliuk, V., and Wilson, D.N. (2018). Structural basis for antibiotic resistance mediated by the Bacillus subtilis ABCF ATPase VmlR. Proc Natl Acad Sci U S A 115, 8978–8983.

Crowe-McAuliffe, C., Murina, V., Turnbull, K.J., Kasari, M., Mohamad, M., Polte, C., Takada, H., Vaitkevicius, K., Johansson, J., Ignatova, Z., et al. (2021). Structural basis of ABCF-mediated resistance to pleuromutilin, lincosamide, and streptogramin A antibiotics in Gram-positive pathogens. Nat Commun 12, 3577.

Cue, D., Lei, M.G., Luong, T.T., Kuechenmeister, L., Dunman, P.M., O’Donnell, S., Rowe, S., O’Gara, J.P., and Lee, C.Y. (2009). Rbf promotes biofilm formation by Staphylococcus aureus via repression of icaR, a negative regulator of icaADBC. J Bacteriol 191, 6363–6373.

Ero, R., Yan, X.F., and Gao, Y.G. (2021). Ribosome Protection Proteins-”New” Players in the Global Arms Race with Antibiotic-Resistant Pathogens. Int J Mol Sci 22.

Fan, R., Li, D., Fessler, A.T., Wu, C., Schwarz, S., and Wang, Y. (2017). Distribution of optrA and cfr in florfenicol-resistant Staphylococcus sciuri of pig origin. Vet Microbiol 210, 43–48.

Fostier, C.R., Monlezun, L., Ousalem, F., Singh, S., Hunt, J.F., and Boel, G. (2021). ABC-F translation factors: from antibiotic resistance to immune response. FEBS Lett 595, 675–706.

Golubev, A., Fatkhullin, B., Khusainov, I., Jenner, L., Gabdulkhakov, A., Validov, S., Yusupova, G., Yusupov, M., and Usachev, K. (2020). Cryo-EM structure of the ribosome functional complex of the human pathogen Staphylococcus aureus at 3.2 A resolution. FEBS Lett 594, 3551–3567.

Graf, M., Huter, P., Maracci, C., Peterek, M., Rodnina, M.V., and Wilson, D.N. (2018). Visualization of translation termination intermediates trapped by the Apidaecin 137 peptide during RF3-mediated recycling of RF1. Nat Commun 9, 3053.

Heymann, J.B. (2018). Guidelines for using Bsoft for high resolution reconstruction and validation of biomolecular structures from electron micrographs. Protein Sci 27, 159–171.

Kasari, V., Pochopien, A.A., Margus, T., Murina, V., Turnbull, K., Zhou, Y., Nissan, T., Graf, M., Novacek, J., Atkinson, G.C., et al. (2019). A role for the Saccharomyces cerevisiae ABCF protein New1 in translation termination/recycling. Nucleic Acids Res. 47, 8807–8820.

Katoh, K., and Standley, D.M. (2013). MAFFT multiple sequence alignment software version 7: improvements in performance and usability. Mol Biol Evol 30, 772–780.

Larsson, A. (2014). AliView: a fast and lightweight alignment viewer and editor for large datasets. Bioinformatics 30, 3276–3278.

Lazaris, A., Coleman, D.C., Kearns, A.M., Pichon, B., Kinnevey, P.M., Earls, M.R., Boyle, B., O’Connell, B., Brennan, G.I., and Shore, A.C. (2017). Novel multiresistance cfr plasmids in linezolid-resistant methicillin-resistant Staphylococcus epidermidis and vancomycin-resistant Enterococcus faecium (VRE) from a hospital outbreak: co-location of cfr and optrA in VRE. J Antimicrob Chemother 72, 3252–3257.

Li, D., Wang, Y., Schwarz, S., Cai, J., Fan, R., Li, J., Fessler, A.T., Zhang, R., Wu, C., and Shen, J. (2016). Co-location of the oxazolidinone resistance genes optrA and cfr on a multiresistance plasmid from Staphylococcus sciuri. J Antimicrob Chemother 71, 1474–1478.

Liebschner, D., Afonine, P.V., Baker, M.L., Bunkóczi, G., Chen, V.B., Croll, T.I., Hintze, B., Hung, L.-W., Jain, S., and McCoy, A.J. (2019). Macromolecular structure determination using X-rays, neutrons and electrons: recent developments in Phenix. Acta Crystallogr D Struct Biol 75, 861–877.

Liu, D., Yang, D., Liu, X., Li, X., Fessler, A.T., Shen, Z., Shen, J., Schwarz, S., and Wang, Y. (2020). Detection of the enterococcal oxazolidinone/phenicol resistance gene optrA in Campylobacter coli. Vet Microbiol 246, 108731.

Lubelski, J., Konings, W.N., and Driessen, A.J. (2007). Distribution and physiology of ABC-type transporters contributing to multidrug resistance in bacteria. Microbiol Mol Biol Rev 71, 463–476.

Marks, J., Kannan, K., Roncase, E.J., Klepacki, D., Kefi, A., Orelle, C., Vazquez-Laslop, N., and Mankin, A.S. (2016). Context-specific inhibition of translation by ribosomal antibiotics targeting the peptidyl transferase center. Proc Natl Acad Sci U S A 113, 12150–12155.

Matzov, D., Eyal, Z., Benhamou, R.I., Shalev-Benami, M., Halfon, Y., Krupkin, M., Zimmerman, E., Rozenberg, H., Bashan, A., Fridman, M., et al. (2017). Structural insights of lincosamides targeting the ribosome of Staphylococcus aureus. Nucleic Acids Res. 45, 10284–10292.

Miller, M.A., Pfeiffer, W., and Schwartz, T. (2010). Creating the CIPRES Science Gateway for Inference of Large Phylogenetic Trees. In Gateway Computing Environments Workshop (GCE) (New Orleans, LA), pp. 1–8.

Minh, B.Q., Schmidt, H.A., Chernomor, O., Schrempf, D., Woodhams, M.D., von Haeseler, A., and Lanfear, R. (2020). IQ-TREE 2: New Models and Efficient Methods for Phylogenetic Inference in the Genomic Era. Mol Biol Evol 37, 1530–1534.

Morroni, G., Brenciani, A., Antonelli, A., D’Andrea, M.M., Di Pilato, V., Fioriti, S., Mingoia, M., Vignaroli, C., Cirioni, O., Biavasco, F., et al. (2018). Characterization of a Multiresistance Plasmid Carrying the optrA and cfr Resistance Genes From an Enterococcus faecium Clinical Isolate. Front Microbiol 9, 2189.

Murina, V., Kasari, M., Hauryliuk, V., and Atkinson, G.C. (2018). Antibiotic resistance ABCF proteins reset the peptidyl transferase centre of the ribosome to counter translational arrest. Nucleic Acids Res. 46, 3753–3763.

Murina, V., Kasari, M., Takada, H., Hinnu, M., Saha, C.K., Grimshaw, J.W., Seki, T., Reith, M., Putrins, M., Tenson, T., et al. (2019). ABCF ATPases Involved in Protein Synthesis, Ribosome Assembly and Antibiotic Resistance: Structural and Functional Diversification across the Tree of Life. J. Mol. Biol.

Murphy, E.L., Singh, K.V., Avila, B., Kleffmann, T., Gregory, S.T., Murray, B.E., Krause, K.L., Khayat, R., and Jogl, G. (2020). Cryo-electron microscopy structure of the 70S ribosome from Enterococcus faecalis. Sci Rep 10, 16301.

Nakane, T., Kimanius, D., Lindahl, E., and Scheres, S.H. (2018). Characterisation of molecular motions in cryo-EM single-particle data by multi-body refinement in RELION. Elife 7.

Orelle, C., Mathieu, K., and Jault, J.M. (2019). Multidrug ABC transporters in bacteria. Res Microbiol 170, 381–391.

Ousalem, F., Singh, S., Chesneau, O., Hunt, J.F., and Boel, G. (2019). ABC-F proteins in mRNA translation and antibiotic resistance. Res Microbiol 170, 435–447.

Pang, S., Boan, P., Lee, T., Gangatharan, S., Tan, S.J., Daley, D., Lee, Y.T., and Coombs, G.W. (2020). Linezolid-resistant ST872 Enterococcus faecium harbouring optrA and cfr (D) oxazolidinone resistance genes. Int J Antimicrob Agents 55, 105831.

Pettersen, E.F., Goddard, T.D., Huang, C.C., Meng, E.C., Couch, G.S., Croll, T.I., Morris, J.H., and Ferrin, T.E. (2021). UCSF ChimeraX: Structure visualization for researchers, educators, and developers. Protein Science 30, 70–82.

Rohou, A., and Grigorieff, N. (2015). CTFFIND4: Fast and accurate defocus estimation from electron micrographs. J Struct Biol 192, 216–221.

Rozov, A., Demeshkina, N., Khusainov, I., Westhof, E., Yusupov, M., and Yusupova, G. (2016). Novel base-pairing interactions at the tRNA wobble position crucial for accurate reading of the genetic code. Nat Commun 7, 10457.

Scheres, S.H., and Chen, S. (2012). Prevention of overfitting in cryo-EM structure determination. Nat Methods 9, 853–854.

Schwarz, S., Zhang, W., Du, X.D., Kruger, H., Fessler, A.T., Ma, S., Zhu, Y., Wu, C., Shen, J., and Wang, Y. (2021). Mobile Oxazolidinone Resistance Genes in Gram-Positive and Gram-Negative Bacteria. Clin Microbiol Rev, e0018820.

Sharkey, L.K., Edwards, T.A., and O’Neill, A.J. (2016). ABC-F Proteins Mediate Antibiotic Resistance through Ribosomal Protection. MBio 7, e01975.

Sharkey, L.K.R., and O’Neill, A.J. (2018). Antibiotic Resistance ABC-F Proteins: Bringing Target Protection into the Limelight. ACS Infect Dis 4, 239–246.

Singh, K.V., Weinstock, G.M., and Murray, B.E. (2002). An Enterococcus faecalis ABC homologue (Lsa) is required for the resistance of this species to clindamycin and quinupristin-dalfopristin. Antimicrob Agents Chemother 46, 1845–1850.

Sivertsen, A., Janice, J., Pedersen, T., Wagner, T.M., Hegstad, J., and Hegstad, K. (2018). The Enterococcus Cassette Chromosome, a Genomic Variation Enabler in Enterococci. mSphere 3.

Su, W., Kumar, V., Ding, Y., Ero, R., Serra, A., Lee, B.S.T., Wong, A.S.W., Shi, J., Sze, S.K., Yang, L., et al. (2018). Ribosome protection by antibiotic resistance ATP-binding cassette protein. Proc Natl Acad Sci U S A 115, 5157–5162.

Svetlov, M.S., Plessa, E., Chen, C.W., Bougas, A., Krokidis, M.G., Dinos, G.P., and Polikanov, Y.S. (2019). High-resolution crystal structures of ribosome-bound chloramphenicol and erythromycin provide the ultimate basis for their competition. RNA 25, 600–606.

Syroegin, E.A., Flemmich, L., Klepacki, D., Vazquez-Laslop, N., Mankin, A.S., Micura, R., and Polikanov, Y.S. (2021). Structural basis for the context-specific action of a classic peptidyl transferase inhibitor. Biorxiv.

Tang, Y., Lai, Y., Wang, X., Lei, C., Li, C., Kong, L., Wang, Y., and Wang, H. (2020). Novel insertion sequence ISChh1-like mediating acquisition of optrA gene in foodborne pathogen Campylobacter coli of swine origin. Vet Microbiol 252, 108934.

Tenson, T., and Mankin, A. (2001). Short peptides conferring resistance to macrolide antibiotics. Peptides 22, 1661–1668.

Tishchenko, S., Gabdulkhakov, A., Nevskaya, N., Sarskikh, A., Kostareva, O., Nikonova, E., Sycheva, A., Moshkovskii, S., Garber, M., and Nikonov, S. (2012). High-resolution crystal structure of the isolated ribosomal L1 stalk. Acta Crystallogr D Biol Crystallogr 68, 1051–1057.

Vazquez-Laslop, N., and Mankin, A.S. (2018). Context-Specific Action of Ribosomal Antibiotics. Annu Rev Microbiol 72, 185–207.

Vorobieva, V., Roer, L., Justesen, U.S., Hansen, F., Frimodt-Moller, N., Hasman, H., and Hammerum, A.M. (2017). Detection of the optrA gene in a clinical ST16 Enterococcus faecalis isolate in Denmark. J Glob Antimicrob Resist 10, 12–13.

Wagner, T., Merino, F., Stabrin, M., Moriya, T., Antoni, C., Apelbaum, A., Hagel, P., Sitsel, O., Raisch, T., Prumbaum, D., et al. (2019). SPHIRE-crYOLO is a fast and accurate fully automated particle picker for cryo-EM. Commun Biol 2, 218.

Wang, Y., Lv, Y., Cai, J., Schwarz, S., Cui, L., Hu, Z., Zhang, R., Li, J., Zhao, Q., He, T., et al. (2015). A novel gene, optrA, that confers transferable resistance to oxazolidinones and phenicols and its presence in Enterococcus faecalis and Enterococcus faecium of human and animal origin. J Antimicrob Chemother 70, 2182–2190.

Waterhouse, A., Bertoni, M., Bienert, S., Studer, G., Tauriello, G., Gumienny, R., Heer, F.T., de Beer, T.A.P., Rempfer, C., Bordoli, L., et al. (2018). SWISS-MODEL: homology modelling of protein structures and complexes. Nucleic Acids Res. 46, W296–W303.

Waterhouse, A.M., Procter, J.B., Martin, D.M., Clamp, M., and Barton, G.J. (2009). Jalview Version 2--a multiple sequence alignment editor and analysis workbench. Bioinformatics 25, 1189–1191.

Watson, Z.L., Ward, F.R., Meheust, R., Ad, O., Schepartz, A., Banfield, J.F., and Cate, J.H. (2020). Structure of the bacterial ribosome at 2 A resolution. Elife 9.

Weaver, K.E., Chen, Y., Miiller, E.M., Johnson, J.N., Dangler, A.A., Manias, D.A., Clem, A.M., Schjodt, D.J., and Dunny, G.M. (2017). Examination of Enterococcus faecalis Toxin-Antitoxin System Toxin Fst Function Utilizing a Pheromone-Inducible Expression Vector with Tight Repression and Broad Dynamic Range. J Bacteriol 199.

Wilson, D.N. (2014). Ribosome-targeting antibiotics and bacterial resistance mechanisms. Nat. Rev. Microbiol. 12, 35–48.

Wilson, D.N., Hauryliuk, V., Atkinson, G.C., and O’Neill, A.J. (2020). Target protection as a key antibiotic resistance mechanism. Nat Rev Microbiol 18, 637–648.

Wilson, D.N., Schluenzen, F., Harms, J.M., Starosta, A.L., Connell, S.R., and Fucini, P. (2008). The oxazolidinone antibiotics perturb the ribosomal peptidyl-transferase center and effect tRNA positioning. Proc Natl Acad Sci U S A 105, 13339–13344.

Zheng, S.Q., Palovcak, E., Armache, J.-P., Verba, K.A., Cheng, Y., and Agard, D.A. (2017). MotionCor2: anisotropic correction of beam-induced motion for improved cryo-electron microscopy. Nat Methods 14, 331–332.

Zhong, X., Xiang, H., Wang, T., Zhong, L., Ming, D., Nie, L., Cao, F., Li, B., Cao, J., Mu, D., et al. (2018). A novel inhibitor of the new antibiotic resistance protein OptrA. Chem Biol Drug Des 92, 1458–1467.

Zivanov, J., Nakane, T., Forsberg, B.O., Kimanius, D., Hagen, W.J., Lindahl, E., and Scheres, S.H. (2018). New tools for automated high-resolution cryo-EM structure determination in RELION-3. Elife 7, e42166.

Zivanov, J., Nakane, T., and Scheres, S.H.W. (2019). A Bayesian approach to beam-induced motion correction in cryo-EM single-particle analysis. IUCrJ 6, 5–17.

